# Imaging-Based Quantitative Assessment of Biomolecular Condensates *in vitro* and in Cells

**DOI:** 10.1101/2024.05.22.594518

**Authors:** Tessa Bergsma, Anton Steen, Julia L. Kamenz, Paola Gallardo, Liesbeth M. Veenhoff

## Abstract

The formation of biomolecular condensates contributes to intracellular compartmentalization, and plays an important role in many cellular processes. The characterization of condensates is however challenging, requiring advanced biophysical or biochemical methods that are often less suitable for *in vivo* studies. A particular need for easily accessible yet thorough methods that enable the characterization of condensates across different experimental systems thus remains. To address this, we present PhaseMetrics, a semi-automated FIJI-based image analysis pipeline tailored for quantifying particle properties from microscopy data. Tested using the FG-domain of yeast nucleoporin Nup100, PhaseMetrics accurately assesses particle properties across diverse experimental setups, including *in vitro*, *Xenopus* egg extracts, and cellular systems. It reliably detects changes induced by various conditions such as the presence of polyethylene glycol, 1,6-hexanediol, a salt gradient, and the molecular chaperone DNAJB6b. By enabling the accurate representation of the variability within the population and the detection of subtle changes at the single particle level, the method complements conventional biochemical assays. Combined, PhaseMetrics is an easily accessible, customizable pipeline that enables imaging-based quantitative assessment of biomolecular condensates *in vitro* and in cells, providing a valuable addition to the current toolbox.

## Introduction

Biomolecular condensates are membraneless compartments that spatially concentrate specific types of biomolecules and are recognized as an important means to, amongst others, sequester cellular components to prevent unwanted interactions, regulate biochemical reactions, and to provide cells with a mechanism to respond to stressors in a fast and adaptive manner (1–5). The formation of biomolecular condensates has generated much excitement and revived an interest in the underlying physicochemical principles of self-assembly processes (1, 2, 6–8). Condensates can form through different physical mechanisms (9, 10). One of these mechanisms includes phase separation, in which once the solubility limit or threshold concentration has been reached, a homogeneous solution separates into a dilute and a dense phase to minimize the overall free energy of the system (1, 2, 10, 11). Despite the extensive progress in probing the driving forces, dynamics and functional implications of biomolecular condensates, a consensus has yet to be reached on the exact combination of experiments necessary to establish whether a condensate can be considered a phase separated compartment or whether distinct principles might have been at play in its formation (12–15). In many instances intrinsically disordered proteins (IDPs) and regions (IDRs) are present in condensates and are implicated in the process of phase separation (1, 9, 16–19).

IDPs encompass a class of proteins that lack a stable three-dimensional structure, and instead exist as a highly dynamic collection of conformations that allow them to perform a wide range of functions in the cell. The dynamic structure of these proteins is largely determined by their amino acid sequences, typically characterized by low-complexity, multivalency, a low proportion of bulky hydrophobic amino acids and a high content of charged and hydrophilic residues (9, 11, 18). Another typical feature of IDPs is their arrangement into stickers-and-spacers, in which motifs with strong interaction potentials (stickers) are separated by motifs with weak interaction potentials (spacers), and the pairwise interaction strength, frequency and relative positioning of these stickers and spacers dictate the saturation concentration for condensate formation (9, 20–23). These features enable them to participate in multivalent homo- and heterotypic protein–protein interactions that define the formation and material properties of condensates (21, 24). Depending on the nature of the interactions, with possible contributions from hydrogen bonding, electrostatic, cation-π, π-π and hydrophobic interactions, different types of assemblies can form ranging from liquid-like to gel-like condensates and solid assemblies. These different phases exhibit distinct physicochemical properties, such as viscosity, composition, and surface tension, which influence their functionality (**Table 1**) (7, 14, 21, 25–27).

**Table 1.** Description of different particle characteristics. Overview of typical characteristics of different types of particles, and techniques that are commonly used to assay their properties.

|  | **Soluble** | **Liquid-like condensates** | **Gel-like condensates** | **Solid-like aggregates** | **Assays** |
| --- | --- | --- | --- | --- | --- |
| *Physical* | Highly mobile | High mobility<br>Fusion and fission | Lower mobility<br>No fusion and fission | Immobile<br>No fusion and fission | FRAP<sup>1)</sup>, timelapse microscopy (fusion and fission, wetting) |
| *Chemical* | - | 1,6-HD and SDS-soluble | Resistance to 1,6-HD, SDS-soluble | 1,6-HD and SDS-insoluble | Sedimentation & turbidity assays, FTA, ThT staining<sup>2)</sup>, fluorescence microscopy |
| *Visual appearance* | Diffuse, homogenous signal | Spherical droplets:<br>- High Circularity<br>- Compact shape<br>- Low density<br>- Homogeneous signal distribution | Denser, irregular particles:<br>- Lower circularity<br>- Less compact shape<br>- Medium/high density<br>- Heterogeneous signal distribution | Fibre-like morphology:<br>- Low circularity<br>- Irregular shape<br>- High density<br>- Heterogeneous signal distribution | Quantitative imaging-based characterization of morphological features |

<sup>1)</sup>FRAP (fluorescence recovery after photobleaching; <sup>2)</sup>ThT (Thioflavin T staining)

One example of a protein complex that heavily relies on IDPs for both its structure and function is the nuclear pore complex (NPC). The NPC is a large protein complex, composed of over 30 types of proteins called nucleoporins (Nups) that each are present in multiple copies. The NPCs regulate the selective transport of macromolecules between the nucleus and the cytoplasm (28–31). Its central channel is composed of intrinsically disordered Nups that are characterized by an enrichment in phenylalanine-glycine (FG) repeats (32–36). *In vitro* studies performed during the last decades have revealed that FG-Nups readily phase separate into liquid-, gel- and amyloid-like particles (33, 37–41), and similarly, when not anchored to the NPC (e.g. prior to assembly or in disease conditions affecting their proper localization), FG-Nups are prone to form condensates and aggregates *in vivo* (42–48). While some of these cytosolic condensates appear to be functional, e.g. serving as assembly platforms for new NPCs (43), surveillance is required to prevent undesirable phase transitions that would interfere with their functionality and proper incorporation into the pore. Mechanisms for the surveillance of FG-Nups (42, 49–53), as well as other IDPs (54–59), are only beginning to be revealed, and a deeper understanding of what drives and guards the transitions between the different possible phase states is lacking. Thus, there is a growing need for experimental tools that enable studying condensate formation and properties in both *in vitro* and *in vivo* settings to unravel the mechanisms and the functional implications of phase state transitions.

Currently, numerous different techniques are being employed to study biomolecular condensates. They include microscopy-based techniques reporting on morphology, and temporal and spatial dynamics, chemical and biophysical approaches assessing features such as reversibility, composition and material properties, and carefully designed mutation and truncation experiments instructed by theory and computational modelling to identify the domains and motifs critical for condensate formation and function (12–15, 20, 27, 60–63). Many of the techniques enabling the assessment of condensate properties, such as Atomic force microscopy (AFM), optical tweezers, fluorescence correlation spectroscopy (FCS), (in-cell) NMR, or microrheology are not widely available and/or require specialized skills and knowledge (12, 13, 27, 63–68).

The simpler imaging or biochemical fractionation-based assays mentioned in **Table 1** are more widely-accessible and commonly used to reveal characteristics of the condensates that enable qualifying them as more liquid-like, gel-like or solid. Biochemical fractionation methods, such as the sedimentation, turbidity and filter trap (FTA) assays, are valuable because they provide quantitative and easy to understand readouts. Yet, considering the widespread heterogeneity in condensate formation, the ability to report on properties at the single condensate level is of particular importance. The main benefit of a microscopy-based method is that it allows monitoring the condensates with spatiotemporal resolution, reports on both morphological and physicochemical properties at single condensate level, and can be used with the same ease across different *in vitro, in cell* and *in vivo* systems. One important concern that has been raised is that many imaging studies rely on descriptive, phenotypic features (13), stressing the need for more detailed, quantitative visual investigations of morphological and physicochemical properties that could assist in the critical assessment of changing material properties of condensates. Several tools have been developed for the quantitative analysis of imaging data (69–72). However, most of these tools were designed for specific applications, and thus are not specifically suited for the analysis of biomolecular condensate properties.

We anticipate that developing a broadly accessible pipeline for obtaining a detailed quantitative description of particle properties from microscopy data could provide valuable insights, particularly when comparing how these properties change with altered experimental conditions. Using the FG-domain of the yeast nucleoporin Nup100 (Nup100FG) as an example case, we demonstrate that the semi-automated FIJI-based image analysis pipeline that we present here, PhaseMetrics, can reliably report on changes in particle properties in response to alterations in the specific experimental conditions and the presence of a specific FG-Nup phase state modulating protein. Furthermore, by testing its performance both *in vitro,* in *Xenopus* egg extracts and in yeast cells, we showcase that the pipeline can be applied with the same ease for both *in vitro* and cellular applications. We validated the output of the image analysis pipeline using commonly-used biochemical assays for studying phase separation. While we only showcase its application for Nup100FG particles, we expect that this versatile pipeline will be equally applicable to any other condensate-forming system.

## Results

### Quantitative imaging-based assessment of crowding effects on Nup100FG particles in vitro

To evaluate the performance of PhaseMetrics, we utilized the FG-domain of the yeast nucleoporin Nup100. The FG-region of yeast Nup100 is highly cohesive and readily undergoes phase separation (41, 73). This tendency can be further accelerated by the use of molecular crowding agents, which are commonly used to mimic the crowded intracellular environment (39–41, 52, 74). Here, we evaluated if PhaseMetrics can quantitatively describe the impact of the crowding agent polyethylene glycol (PEG) on Nup100FG particle properties (**Figure 1**), and if the quantifications align with our previous biochemical characterizations of the particles (42, 52). Of note is that while PEG is generally considered an inert molecule (11, 74), for some macromolecules it instead promotes phase separation through co-condensation (74–76). This also appears to apply for Nup100FG, as PEG particles are observed in the presence of Nup100FG (**Figure S3A-C**). Thus, the effect of PEG in our experiments cannot be considered a true crowding effect; instead it appears to reflect associative phase separation (74, 76). In a physiological salt buffer (100 mM Tris, 150 mM NaCl, pH 8.0), Nup100FG forms irregularly shaped structures of varying sizes, while in the presence of 10% PEG3350, we observe a more homogeneous particle population of brighter and more circular particles (**Figure 1A, Figure S1A,D**). Using the PhaseMetrics pipeline we quantitatively assessed 300 Nup100FG particles formed both in the absence and presence of 10% PEG3350, and find that in its presence, the particles mean intensity, size, perimeter, and circularity increase significantly (**Figure 1B,D-F**). Moreover, in the presence of a crowding agent, the intensity within the particles is often less variable (less particles have skewness values deviating from 0) (**Figure 1C**). In general, the particle population is more homogeneous in the presence of 10% PEG3350 as reflected by the Nup100FG particles becoming more uniform in skewness, size, perimeter and circularly (**Figure 1C-F**). The difference in intensity of the particles formed in the presence or absence of PEG3350 merely indicates that the particles are different, as a direct interpretation in terms of concentration of Nup100FG protein is not possible as quenching effects may play a significant role. The particles shown in **Figure 1A** were selected using a pre-designed excel sheet for the selection of the particle that is most representative for the overall particle population, based on all analysis parameters (**Figure 1B-F**). This also holds true for the representative particles shown in the remaining figures of this study. To allow for a measurement of the soluble fraction, we performed an identical experiment in the droplet-in-chamber setup (**Figure 8B, Figure S1D**), which revealed that less protein remains in the dilute phase in the presence of 10% PEG3350 (**Figure 1G**), leading to a higher partition coefficient (**Figure 1H**).

**Fig. 1.**
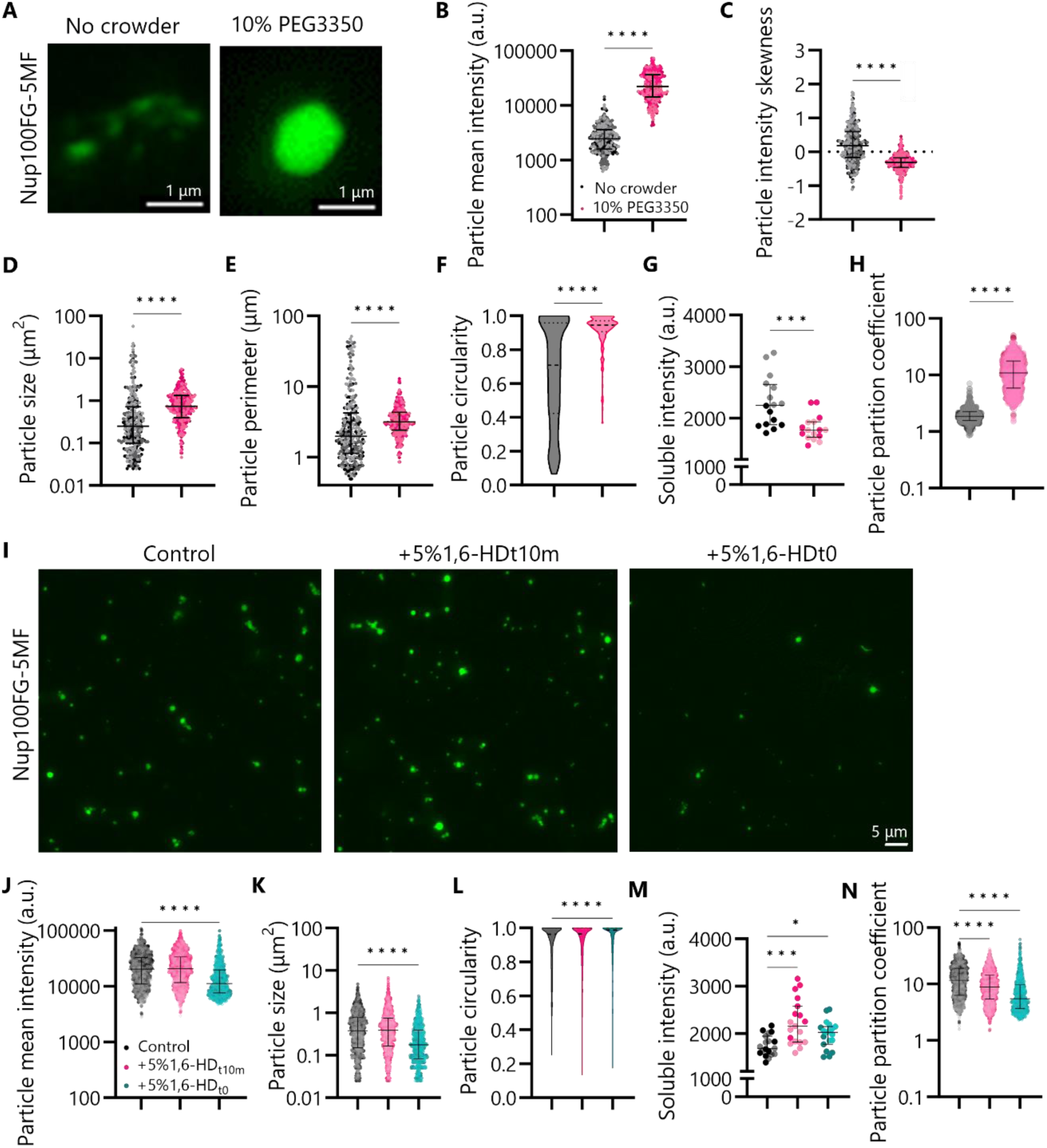
Quantitative characterization of the chemical properties of Nup100FG particles formed in vitro. **(A)** Representative images of Nup100FG-5MF particles, formed in the absence or presence of 10% PEG3350 (1h) (glass-slide setup). The images are selected using the pre-designed excel sheet for the selection of the particle that is most representative for the overall particle population, based on the analysis parameters quantified in B, C, D, E, F. Scale bar, 1 µm. **(B-F)** Mean fluorescence intensity, intensity skewness, size, perimeter and circularity of Nup100FG-5MF particles exemplified in **(A)**. Graphs show median ± interquartile range of 300 particles per condition (n=3). 100 particles were analysed for each independent replicate. ***P<0.001, ****P<0.0001. **(G,H)** Mean intensity of soluble fraction and partition coefficient of Nup100FG-5MF protein mixtures (droplet-in-chamber setup; particles exemplified in **Figure S1D**). **(I)** Representative images of Nup100FG-5MF particles, formed in the presence of 10% PEG3350 (1h) (droplet-in-chamber setup), in the absence or upon exposure to 5% 1,6-HD, for either 10 minutes after an hour of particle formation (t10m), or immediately from the start (t0). Scale bar, 5 µm. **(J-L)** Mean fluorescence intensity, size and circularity of Nup100FG-5MF particles exemplified in **(I)**. Graphs show median ± interquartile range of ≥750 particles per condition (n=2). **(M,N)** Mean intensity of soluble fraction and partition coefficient of Nup100FG-5MF protein mixtures exemplified in **(I)**. Graph shows median ± interquartile range of 19 images per condition (n=2). **P<0.01, ****P<0.0001.

The increase in particle mean intensity (**Figure 1B**) and size (**Figure 1D**), and decrease in the soluble fraction (**Figure 1G**) observed in the presence of 10% PEG3350 align with previous data from sedimentation assays (42, 52). These assays revealed that in the absence of a crowding agent, Nup100FG mostly remains in the soluble fraction (42), whereas in the presence of 10% PEG3350 most of the protein re-distributes to the dense, phase separated fraction (52). Similarly, filter trap assays (FTA) revealed that in the presence of 10% PEG3350 more protein gets trapped on the filter trap membrane than in the absence of a crowding agent (52). Combined, the PEG-induced changes quantified using the image-based analysis align with those previously reported by us using biochemical methods (42, 52).

### Quantitative assessment of the sensitivity of Nup100FG particles to 1,6-HD

The intermolecular interactions that drive the formation of the distinct phases also define the sensitivity of the particles to changes in their surrounding environment. Liquid-like condensates, for example, are characterized by weak intermolecular interactions, and generally are more sensitive to external perturbations (27). The aliphatic alcohol 1,6-hexanediol (1,6-HD) can perturb weak hydrophobic interactions, and is commonly used as a tool to assess the liquidity of condensates (14, 60, 77), whereas lower 1,6-HD concentrations mainly dissolve liquid-like condensates, higher 1,6-HD concentrations have been reported to also dissolve gel-like condensates (27, 78). PhaseMetrics was used to assess the effect of 1,6-HD on the Nup100FG particles formed in the presence of 10% PEG3350 (using the droplet-in-chamber setup). Three conditions were compared (**Figure 1I**), one in which the Nup100FG particles were not exposed to 1,6-HD, one in which preformed (1 hour) Nup100FG particles were exposed for 10 minutes to 5% 1,6-HD (1,6-HD_t10m_), and one in which the Nup100FG particles were formed (1 hour) in the presence of 5% 1,6-HD (1,6-HD_t0_). The quantitative assessment of the effect of 1,6-HD treatment on preformed Nup100FG particles mainly served to assess the fraction of liquid-like condensates that remained in the mixture after one hour. It revealed no significant changes in particle intensity, size, and circularity (**Figure 1J-L**), suggesting that the main population of Nup100FG particles is either gel- or solid-like. This is consistent with what has previously been published (41). While the majority of protein is in 1,6-HD-resistent particles, the observed increase in the soluble fraction under these conditions (**Figure 1M**) suggests that a small fraction of Nup100FG is solubilized by 1,6-HD. The formation of Nup100FG particles in the presence of 1,6-HD served to show the contribution of weak hydrophobic interactions to Nup100FG phase separation. In this condition, less particles are formed (**Figure 1I, Figure S1F**), and the observed particles are dimmer (**Figure 1J**) and smaller (**Figure 1K**). Consistently, a larger proportion of Nup100FG remains in the dilute, soluble fraction (**Figure 1M**). Moreover, in line with the soluble measurements, a drop in the partition coefficient was observed for both conditions (**Figure 1N**). In sum, we conclude that the imaging-based assessment of Nup100FG particles provides a means to quantify the impact of 1,6-HD on particle properties, and the analysis supports that the majority of particles are formed through weak hydrophobic interactions and are gel- or solid-like after 1 hour.

### Quantitative assessment of particle formation and properties in response to altering ionic strength

We next investigated how Nup100FG particle properties are affected by alterations in the ionic strength of the buffer solution. For this purpose, we made use of increasing concentrations of sodium chloride (NaCl), and compared particle formation in the absence or presence of 10% PEG3350. In the absence of a crowding agent (**Figure 2**), amorphous structures are observed when the particles are formed at zero or 150 mM NaCl. As the concentration of NaCl increases, however, a transition is observed from irregularly shaped structures towards circular species. Overall, with increasing salt concentration, particle mean intensity, size, perimeter and circularity increase, and the skewness decreases (**Figure 2B-G**). The signal distribution within the particles becomes more uniform, and the distribution of particle perimeters becomes more homogeneous, indicative of the structures becoming more homogeneous and compact.

**Fig. 2.**
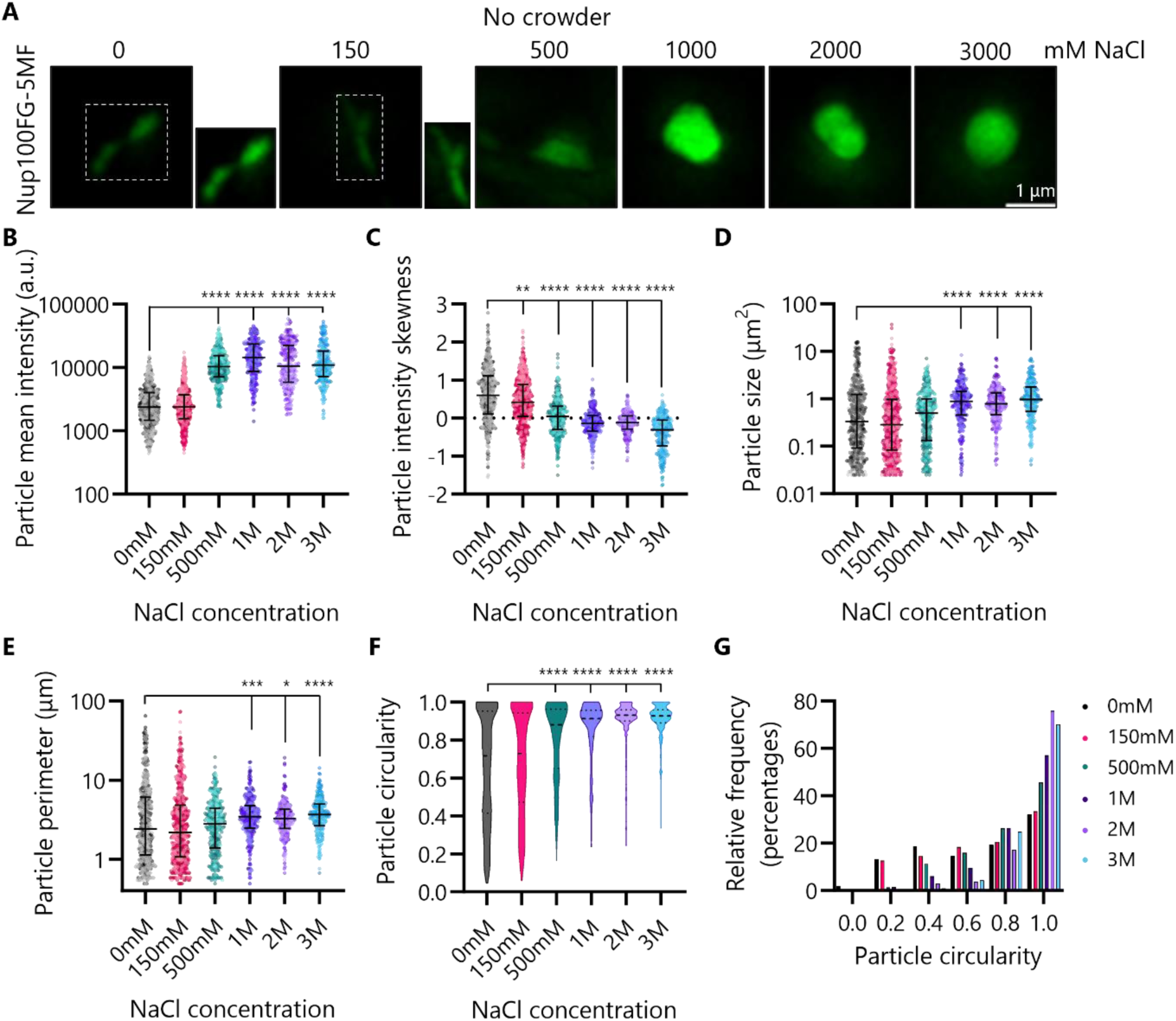
Quantitative assessment of salt effects on Nup100FG particles formed in vitro in absence of a crowding agent. **(A)** Representative images of Nup100FG-5MF particles, formed in the absence of a crowding agent, in response to varying salt concentrations (1h) (glass-slide setup). Insets correspond to images with 2x enhanced brightness, to highlight low-intensity particles. Scale bar, 1 µm. **(B-G)** Mean fluorescence intensity, intensity skewness, size, perimeter, circularity and frequency distribution of the circularity of Nup100FG-5MF particles exemplified in **(A)**. All graphs show median ± interquartile range of ≥225 particles per condition (n=3). ≥75 particles were analysed for each independent replicate. The means of each condition were compared to the mean of the 0mM condition. *P<0.05, **P<0.01, ***P<0.001 ****P<0.0001.

In the presence of 10% PEG3350 (**Figure 3**) Nup100FG forms circular particles even in the absence of salt, but the particles become more compact with increasing salt concentrations. Particle intensity, size and perimeter decrease, and overall, the signal distribution throughout the particles becomes slightly more homogeneous (**Figure 3B-E**). The circularity of the particles does not appear to be strongly affected by the salt (**Figure 3F,G**). The impact of salt on the solubility of proteins has been ascribed to hydrophobic effects (79–83), with in particular salts earlier in the Hofmeister series (including NaCl) strengthening hydrophobic interactions (79). The observation of homotypic particle formation (no requirement of PEG) at high NaCl concentrations suggests that Nup100FG particle formation under these circumstances is stimulated by screening of electrostatic interactions, and strengthening of non-charged, hydrophobic interactions, which is in line with what has been previously published for Nup100 paralogs (73, 84). Interestingly, and aligning well with both PEG and NaCl heightening the hydrophobic effect, we see that the particles that are formed in the absence of crowder at the higher NaCl concentrations, ranging from 1 to 3M, display similar intensity, size and circularity values as those observed in the presence of 10% PEG3350 in the lower salt regime (**Figure 3H-J**). Overall, the results from each the crowder, 1,6-HD and salt experiments, indicate that the pipeline is capable of reporting on changing particle properties in response to alternating environmental conditions, and might aid in the distinguishment between different types of particles.

**Fig. 3.**
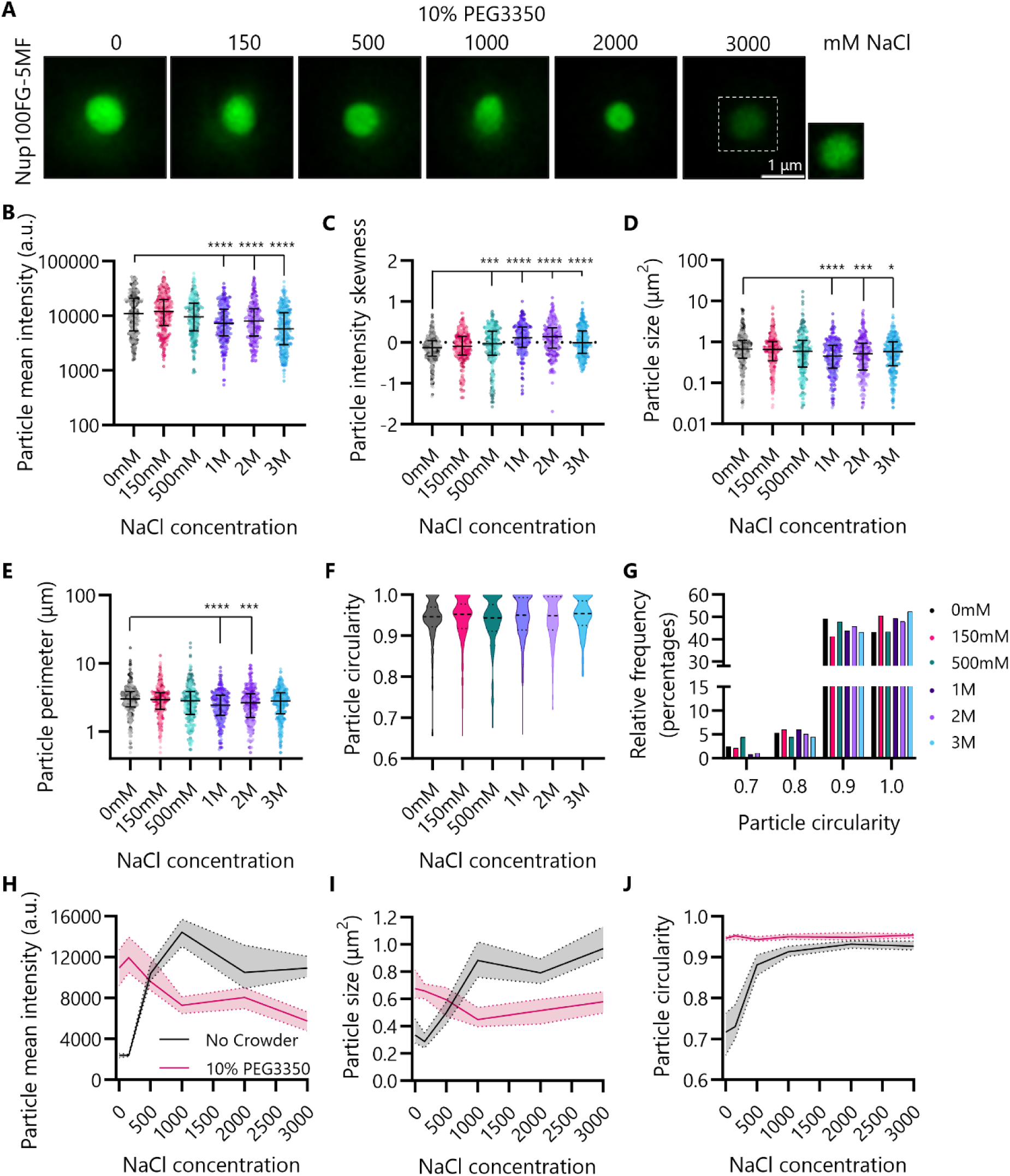
Quantitative assessment of salt effects on Nup100FG particles formed in vitro in presence of a crowding agent. **(A)** Representative images of Nup100FG-5MF particles, formed in the presence of 10% PEG3350, in response to varying salt concentrations (1h) (glass-slide setup). Insets correspond to images with 2x enhanced brightness, to highlight low-intensity particles. Scale bar, 1 µm. **(B-G)** Mean fluorescence intensity, intensity skewness, size, perimeter, circularity and frequency distribution of the circularity of Nup100FG-5MF particles exemplified in **(A)**. All graphs show median ± interquartile range of ≥250 particles per condition (n=3). ≥75 particles were analysed for each independent replicate. The means of each condition were compared to the mean of the 0mM condition. *P<0.05, ***P<0.001, ****P=<0.0001. **(H-J)** Overlapping intensity, size and circularity distributions between particles observed in absence and presence of 10% PEG3350 in the low and high salt regimes. In all graphs, the bold lines plus shades show the median and ± 95% CI of all measurements from three replicates.

### Quantitative assessment of particle properties in the presence of a phase state modulating protein

Previously, we (52) and others (53) identified DNAJB6b as a modulator of FG-Nup phase transitions. Making use of sedimentation assays and FTAs, we showed that DNAJB6b delays the time-dependent transition of FG-Nups into solid-like aggregates and maintains FG-Nups for a longer period in the soluble and liquid-like state (52). To complement these bulk biochemical assays, it is valuable to gain information on the singe particle level. Hence, we tested if the analysis of large numbers of particles using PhaseMetrics was able to detect DNAJB6b-dependent changes in Nup100FG particle properties. For this purpose, we followed Nup100FG particle formation over a prolonged time frame, both in the absence and presence of DNAJB6b (**Figure 4A**). This revealed that Nup100FG particle properties progressively change over time, showing an increase in intensity and size and a drop in particle circularity (**Figure 4B-D, Figure S2A-C**), fitting with a transition into more solid phases. This progression is delayed in the presence of DNAJB6b (1:1 ratio). We plotted the mean fluorescence intensity of Nup100FG particles against the FTA dataset reporting on the SDS-insoluble fraction of Nup100FG that we published previously (52) (**Figure 4E**). This highlighted that the kinetics of the phase state transitions towards more aggregated species are equally well reported by the imaging-based quantification and the FTA. At later time points, the results from the imaging-based read-out and FTA deviate, which may be attributed to incomplete recovery of the SDS-insoluble Nup100FG fraction. Lastly, in the presence of DNAJB6b a larger pool of Nup100FG monomers remains in the dilute phase (**Figure 4G, Figure S2D**) and less protein partitions into particles (**Figure S2E**). These observations are also in line with our previous biochemical data showing that DNAJB6b maintains a larger fraction of Nup100FG in the soluble state (52).

**Fig. 4.**
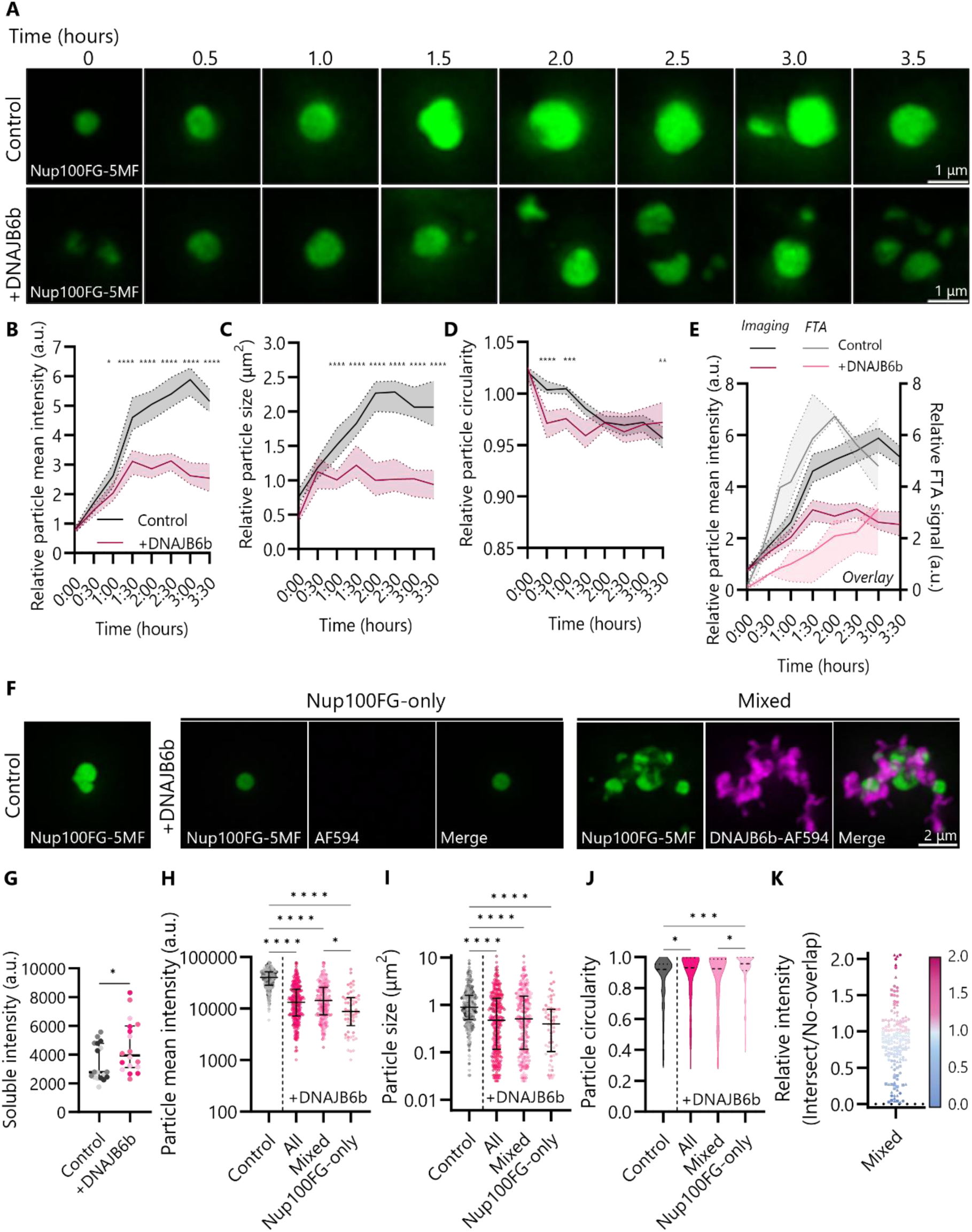
Quantitative assessment of changing Nup100FG particle properties in vitro in the presence of a phase state modulator. **(A)** Representative images showing changing properties of Nup100FG-5MF particles, formed in the presence of 10% PEG3350, over prolonged timeframe in the absence and presence of DNAJB6b (glass-slide setup). Scale bar, 1 µm. **(B-D)** Mean fluorescence intensity, size and circularity of Nup100FG-5MF particles exemplified in **(A)**, relative to the mean of the control at t0. The bold lines plus shades show the median ± 95% CI of all measurements from three replicates. 100 particles were analysed for each time point for each of the independent replicate experiments. *P<0.05, **P<0.01, ***P<0.001, ****P<0.0001. **(E)** Comparison between our previously published FTA dataset^40^ (relative signal intensity to mean control at t15) and relative mean fluorescence intensity of Nup100FG-5MF particles (relative to mean control at t0) formed in the presence of DNAJB6b exemplified in **(A)**. The bold lines plus shades show the median ± 95% CI from three replicates. **(F)** Representative images showing Nup100FG-5MF particles in the absence or presence of DNAJB6b-AF594 (1h) (molar ratio 1:1) (glass-slide setup). Scale bar, 2 µm. **(G)** Mean intensity of soluble fraction of Nup100FG-5MF±DNAJB6b protein mixtures exemplified in **Figure S2D** (droplet-in-chamber setup). Graph shows median ± interquartile range of 18 images per condition (n=3). *P=<0.05. **(H-J)** Mean fluorescence intensity, size and circularity of Nup100FG-5MF particles exemplified in **(F)**. Mixed: Nup100FG particles that are colocalized with a DNAJB6b particle, Nup100FG-only: Nup100FG particles that are not colocalized with a DNAJB6b particle. Graphs show median ± interquartile range of 300 particles per condition (n=3). 100 particles were analysed for each independent replicate. *P<0.05, ***P<0.001, ****P<0.0001. **(K)** Comparison of mean fluorescence intensity in intersected and no-overlap regions of Nup100FG-5MF+DNAJB6b-AF594 particles. Graph shows the relative intensity in the intersect over no-overlap regions for a total of 150 particles (n=3; 50 particles were analysed for each independent replicate). The individual particles were coloured according to their relative intensity values as indicated in the heat map, in which blue reflects a decrease and pink reflects an increase in intensity within the intersected region as compared to the no-overlap region. Different symbols were used to highlight the individual datapoints belonging to each of the independent replicates.

To allow for a more detailed assessment of the impact of DNAJB6b on the properties of Nup100FG particles, we made use of fluorescently-labelled DNAJB6b (**Figure 4F**). Similar to the unlabelled DNAJB6b we observed a strong reduction in particle intensity and size, and an increase in circularity of the particles that are formed in the presence of DNAJB6b (**Figure 4H-J**, **Figure S2F**, ‘all’). In the presence of fluorescently-labelled DNAJB6b we can further discriminate Nup100FG particles that are not co-colocalized with DNAJB6b (Nup100FG-only) and those that are colocalized with a DNAJB6b particle (mixed). Comparison of these subgroups clearly shows that the particles of Nup100FG alone are less intense, smaller and rounder (**Figure 4F,H-J**, **Figure S2F**). This may indicate that DNAJB6b associates more strongly with the more solid-like particles compared to the more liquid-like condensates. Interestingly, a more close evaluation of the particles that are colocalized with a DNAJB6b particle shows that the intensity at the sites of contact between the Nup100FG and DNAJB6b particles is frequently lower than the intensity at regions where there is no overlap (**Figure 4K**, **Figure S2G)**. Overall, the results align well with the biochemical assays, while additionally providing information at the single particle as well as intra-particle level, supporting the applicability of the pipeline to report on changes in particle properties in the presence of a phase state modulator.

### Quantitative assessment of Nup100FG particle formation and properties in Xenopus laevis egg extracts (XEE)

Whilst *in vitro* experiments have proven to be useful in gaining fundamental insights into the driving forces and physicochemical properties of biomolecular condensates, these systems are a strong oversimplification of the complex cellular environment. For this reason, once the fundamental principles have been mapped using simpler *in vitro* systems, a transition towards more complex systems is often made, to confirm whether the same phenomena can be observed under physiologically relevant conditions (12, 14). *Xenopus laevis* egg extracts (XEE) may present a valuable bridge between *in vitro* and cellular experiments by maintaining the flexibility for manipulations that one typically has with *in vitro* systems, yet at the same time providing a complex, crowded, and proteinaceous environment that may better represent the native cytosol. Here, we introduced the same purified Nup100FG fragment as used in our *in vitro* experiments into XEE, and used PhaseMetrics to track Nup100FG particle properties over a three-hour time frame (**Figure 5**). Overall, over time we observed an increase in particle mean intensity, intensity skewness, size and perimeter, and a drop in circularity (**Figure 5B-F**). Other than the much more distinguished drop in the circularity parameter after three hours (**Figure 5F,H**), Nup100FG condensates appear to undergo a similar time-dependent progression towards more gel and/or solid-like species as observed in our *in vitro* experiments with chemically-defined buffers. Together these results indicate that PhaseMetrics is equally suited for detecting changing particle properties in a more complex experimental system.

**Fig. 5.**
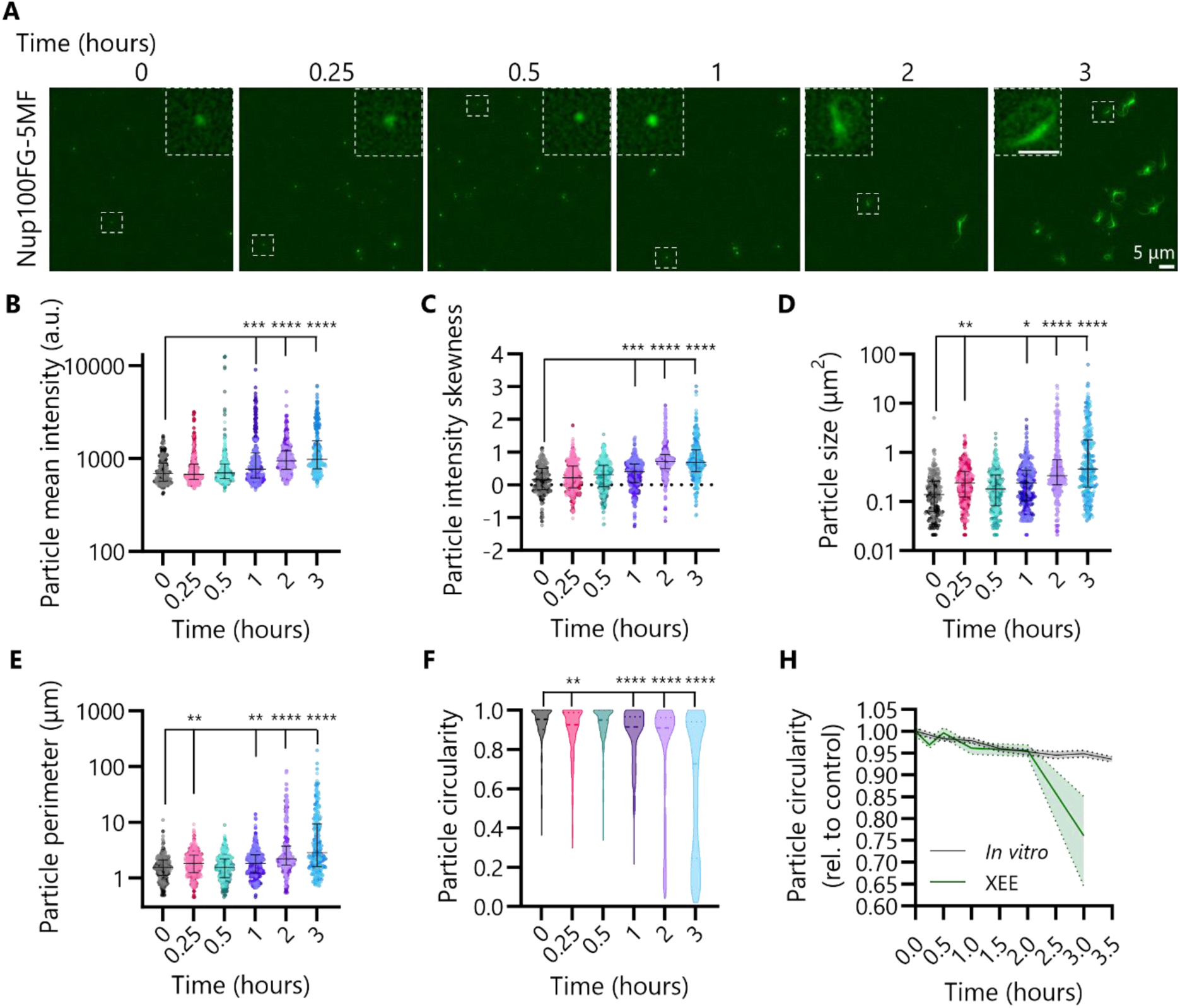
Quantitative assessment of changing Nup100FG particle properties in Xenopus egg extracts over time. **(A)** Representative images of Nup100FG-5MF particles in Xenopus egg extracts (XEE), analysed at the indicated time points. Scale bar, 5 µm. Insets represent Zoom-in panels illustrating the most-representative particle for each time point. Scale bar, 2 µm. **(B-F)** Mean fluorescence intensity, skewness, size, perimeter and circularity of Nup100FG-5MF particles exemplified in **(A)**. Graphs show median ± interquartile range of 300 particles per condition (n=3). The means of each timepoint were compared to the mean of the t0 timepoint. *P<0.05, **P<0.01, ***P<0.001, ****P<0.0001. **(H)** Comparison of time-dependent changes in circularity of Nup100FG-5MF particles exemplified in (**A**) (XEE) and Figure 4A (in vitro, formed in the presence of 10% PEG3350). The bold lines plus shades show the median and ± 95% CI of all measurements from three replicates.

### Quantitative assessment of Nup100FG particle formation and properties in yeast cells

After having established that the pipeline works equally well on Nup100FG particles formed in both an *in vitro* system and in XEE, we tested PhaseMetrics on live yeast cells. For this purpose, we overexpressed eGFP-Nup100FG in *S. cerevisiae* and tracked eGFP-Nup100FG particles over a six-hour timeframe (**Figure 6**). For assessing the performance of the pipeline, we compared the results obtained with a manual quantification. Manually, the cells were categorized as not having particles, having 1, 2, ≥3 more circular particles (subjectively referred to as condensates), having particles with fibre-like protrusions (referred to as amyloid-like bodies (AB)), and having particles that were non-round but also not fibre like (referred to as amorphous aggregates (AA). A time-dependent progressive decrease in the percentage of cells with condensates, concomitantly with an increase of the percentage of cells with no condensates and cells with AB structures is observed (**Figure 6B, Figure S3A**). A similar outcome was obtained with the pipeline-based analysis (**Figure 6C, Figure S3B**). Importantly, PhaseMetrics outperformed the manual analysis by enabling the analysis of much larger numbers of cells in less time and by enabling a further stratification of the three subclasses, to also separate cells containing multiple types of particles. Moreover, it simultaneously allows for the extraction of relevant parameters at the single particle level, showing that particle intensity, size, and perimeter increase, while particle circularity decreases (**Figure 6D-G**), which fits with the predicted transition into more gel-like and solid-like phases.

**Fig. 6.**
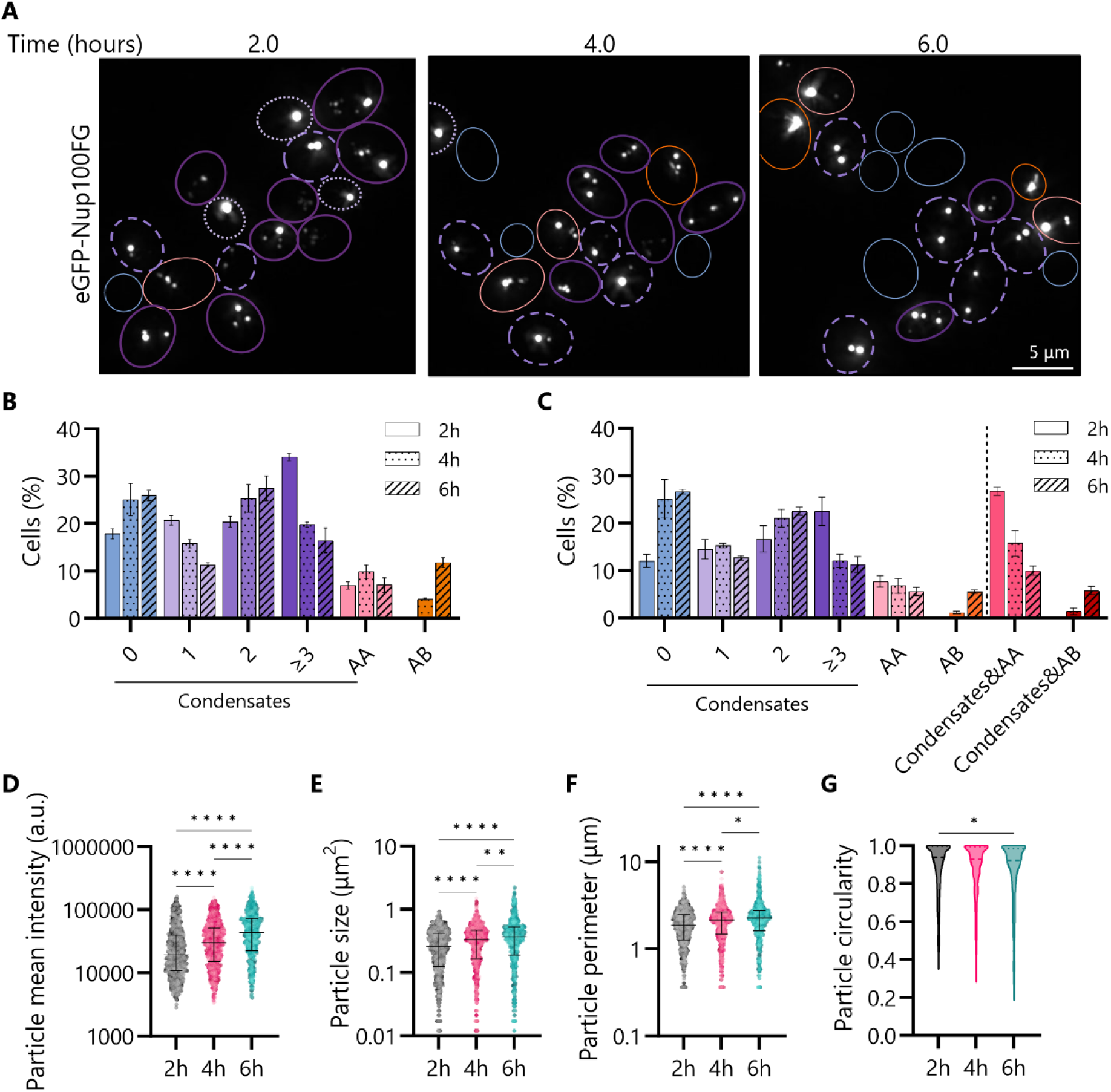
Quantitative assessment of changing Nup100FG particle properties in yeast cells over time. **(A)** Representative images of eGFP-Nup100FG particles in yeast cells, analysed at the indicated timepoints. Cells are outlined with coloured circles aligning with the different subcategories as specified in **(B)**. Scale bar, 5 µm. **(B,C)** Distributions of eGFP-Nup100FG particle subpopulations per cell, using **(B)** manual and **(C)** plugin-based quantification. AB: amyloid-like body, AA: amorphous aggregate. Graphs B-C show mean ± SEM of ≥450 cells per condition (n=3). 200-400 particles were counted for each independent replicate. **(D-G)** Mean fluorescence intensity, size, perimeter and circularity of eGFP-Nup100FG particles exemplified in **(A)**. Graphs D-G show median ± interquartile range of ≥750 particles per condition (n=3). *P<0.05, **P<0.01, ****P<0.0001.

We next set out to assess whether the pipeline also could be used to pick up changes in the properties of the eGFP-Nup100FG particles in the presence of a known FG-Nup phase state modulator. For this purpose, we co-overexpressed eGFP-Nup100FG with mCherry-DNAJB6b (**Figure 7A**). Comparison of the particle distribution between the manual and pipeline-based quantifications again revealed very similar outcomes (**Figure 7B,C, Figure S3C,D**). A clear reduction in the number of AB’s is seen in the presence of DNAJB6b, whereas the remaining subpopulations are not significantly affected (**Figure S3C,D**). From the assessment of the particles properties we also find support for a role of DNAJB6b in favouring the liquid/gel-like phases by preventing their transition towards the more solid phases, as we see that in the presence of DNAJB6 there is a reduction in the number of AB’s and the soluble Nup100FG fraction is larger (**Figure 7G,H**). No changes in the size and perimeter of the eGFP-Nup100FG particles were detected in the presence of DNAJB6b (**Figure 7E,F**), albeit that we see a loss of particles with particularly large perimeters (5.5-10). The difference in intensity of the particles formed in the presence or absence of DNAJB6b (**Figure 7D**) indicates that the particles are different; as mentioned before, a direct interpretation in terms of concentration of Nup100FG protein is not possible as quenching may play a significant role.

**Fig. 7.**
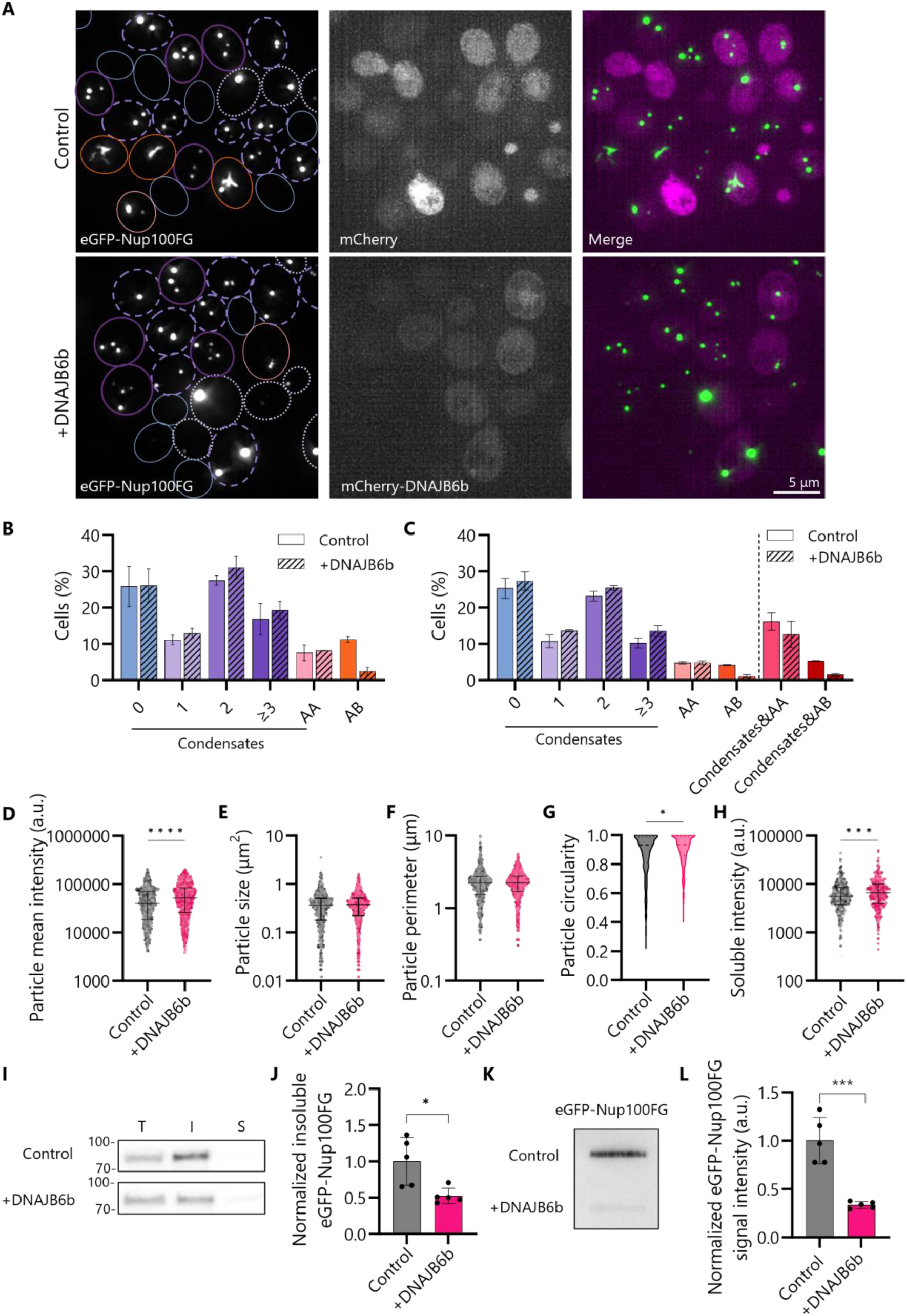
Quantitative assessment of changing Nup100FG particle properties in yeast cells in the presence of a phase state modulator. **(A)** Representative images of eGFP-Nup100FG particles in yeast cells in absence (mCherry as negative control) or presence of mCherry-DNAJB6b. Scale bar, 5 µm. **(B,C)** Distributions of eGFP-Nup100FG particle subpopulations per cell, using **(B)** manual and **(C)** plugin-based quantification. AB: amyloid-like body, AA: amorphous aggregate. Graphs B-C show mean ± SEM of ≥500 cells per condition (n=3). 250-350 particles were counted for each independent replicate. **(D-G)** Mean fluorescence intensity, size, perimeter and circularity of eGFP-Nup100FG particles exemplified in **(A).** *P<0.05, ****P<0.0001. **(H)** Mean intensity of eGFP-Nup100FG soluble fraction. Graphs D-H show median ± interquartile range of ≥800 particles per condition (n=3). ***P<0.001. **(I)** Sedimentation assay to assess the soluble and insoluble fraction of eGFP-Nup100FG in the absence or presence of DNAJB6b. **(J)** Quantification of the insoluble fraction of eGFP-Nup100FG. Represented band intensities are relative to the average intensity of the control. Mean ± SEM (n=5). *P=<0.05. **(K)** Filter trap assay to assess aggregated fraction of eGFP-Nup100FG in the absence or presence of DNAJB6b. **(L)** Quantification of the band intensities of eGFP-Nup100FG on filter trap. Represented band intensities are relative to the average intensity of the control. Mean ± SEM (n=5). ***P<0.001.

To corroborate the results from the imaging read-out suggesting that DNAJB6b delays the transition towards more solid-like species and keeps the protein in a more liquid-like phase, we performed sedimentation assays and FTAs using cell lysate of the yeast cells co-overexpressing eGFP-Nup100FG and mCherry-DNAJB6b (mCherry alone as negative control) (**Figure S4**). Assessment of the soluble and insoluble protein fractions using sedimentation assays, revealed a reduction in the insoluble fraction (**Figure 7I,J**), and trend towards an increased amount of soluble eGFP-Nup100FG in the presence of DNAJB6b (**Figure S4A,B**). The FTA revealed a reduction in the aggregated fraction for eGFP-Nup100FG in the presence of DNAJB6b (**Figure 7K,L, Figure S4C**). These observations are in accordance with the observed increase in soluble intensity and reduction in AB’s in the imaging dataset. Combined, PhaseMetrics is equally applicable for assessment of particle properties in both *in vitro* and cellular applications, and the read-outs obtained using this type of analysis appear complementary to other physicochemical phase separation assays.

## Discussion

By exploring the physicochemical properties of different phase states and employing multidisciplinary approaches, researchers have made significant strides in our overall understanding of biomolecular condensates. However, further advancements require expanding the current toolbox for studying the mechanisms underlying their formation and functional relevance in the cellular context. In particular, there is a need for easily accessible, yet thorough strategies to characterize condensates across different experimental systems. Here, based on the characteristic physicochemical properties and morphological features of different particles, as summarized in **Table 1**, we developed PhaseMetrics to report on a relevant combination of properties to characterize particles, and how these change over time or in response to multiple interventions. We demonstrate that the pipeline functions well across different experimental systems and represents a good complement to conventional biochemical assays. The quantitative analysis of hundreds of particles gives a more honest representation of the variability within particle populations and aids in the unbiased selection of those particles that are most representative. Additionally, the method is sensitive enough to detect minor particle subpopulations and small changes in particle properties, both in response to alternating environmental conditions and the presence of a phase state modulating protein. Whilst liquid-like particles, gel-like particles, and aggregates cannot be discriminated from one another purely based on their morphological features, their response to alternating experimental conditions can give additional insights into their physicochemical nature. Moreover, demonstrating changes in particle properties under biologically relevant conditions, such as the presence of a phase state modulator, a post-transcriptional modification, or a change in pH, may be as relevant as defining the exact nature of the particles.

### PhaseMetrics faithfully reports on known chemical phase state modulators

In the *in vitro* experiments, we specifically selected a set of interventions for which we knew, based on prior experimentation and theory, how they should impact condensation. This allowed us to assess whether the changes in particle properties reported by PhaseMetrics would match the theoretically or experimentally predicted physicochemical properties that define condensates and aggregates. Indeed, PhaseMetrics can be used to detect the anticipated effects for each of the selected conditions, including the addition of PEG, 1,6-HD and a salt gradient, and aid in distinguishing between different types of particles that are formed.

PEG modulates the extent to which a protein interacts with itself (both intra- and intermolecular) and/or with the surrounding solvent (40, 76, 85, 86). Indeed, in the presence of 10% PEG3350 a more homogeneous population of Nup100FG particles with increased intensity, size and roundness was formed. This may indicate that PEG favours the formation of condensates or stabilizes them, at the expense of the competing transitions to amorphous structures. The observed overall reduction in particle abundance and increase in the soluble fraction upon 1,6-HD treatment are in line with previous studies and the reported importance of hydrophobic interactions for Nup100FG particle formation (41, 52). The observed shift towards a more uniform and compact particle population with increasing NaCl concentrations further highlights the importance of hydrophobic interactions.

Whilst in this study we used PEG, 1,6-HD, and salts to modulate the hydrophobic interactions that are important to the formation of Nup100FG particles, similar studies could be performed for other proteins that rely on other interaction types, such as electrostatic, cation-pi or polar interactions. These kinds of studies could aid in better linking the observed changes to the molecular driving forces controlling the formation and material properties of the particles (79, 87). Moreover, thorough mapping of how particles respond to external stimuli can also aid in improving our understanding of the adaptability and stability of condensates in the cellular context.

### PhaseMetrics complements biochemical bulk assays to report on subtle changes at the single particle level

Besides changing buffer conditions, we tested the ability of PhaseMetrics to report on changing particle properties *in vitro* in the presence of DNAJB6b, known for its ability to modulate FG-Nup phase transitions (52, 53). The observed increase in the soluble fraction, increased circularity and altered intensity of the Nup100FG particles in the presence of DNAJB6b, are in line with our previously performed sedimentation assays and FTAs (52), and are indicative of the ability of DNAJB6b to delay the transition towards more solid-like species. The slight increase in the soluble pool could be interpreted as DNAJB6b stabilizing the monomeric species. Additive to the previous biochemical characterizations is the observation that the Nup100FG particles that are adjacent to DNAJB6b particles are more intense, larger, and less round than those that are not colocalizing with DNAJB6b. This may imply that DNAJB6b associates more strongly with the more solid-like particles, whereas it has a more dynamic interaction with the more liquid-like condensates. Observations like this illustrate the importance of performing sensitive large-scale quantitative assessments of particle properties and the potential to direct future studies addressing how DNAJB6b influences FG-Nup phase transitions mechanistically.

### Xenopus egg extracts can serve as a valuable experimental system to study biomolecular condensates

We explored the use of XEE in condensation assays. XEE are known for their ability to recapitulate complicated cellular reactions, while maintaining the flexibility for manipulations that one typically has with *in vitro* systems (12, 14). The characterization of Nup100FG particles in the XEE environment revealed that they shared overlapping features with those observed in the chemically defined buffers. The largest difference between the two experimental setups appears to be the earlier appearance of species with fibre-like protrusions, which could be indicative of a faster progression towards more solid-like phases in the extracts. Possible explanations for this could be differences in the extend of crowding between the two conditions, the presence of native, unknown XEE factors that co-aggregate with Nup100FG or a more direct influence of PEG entering the Nup100FG particles. Also, the extracts provide a much more dynamic environment compared to the *in vitro* environment and this likely also influences the local environment and properties of the observed condensates (12). Overall, the relative fast progression towards amyloid-like bodies in the XEE can be considered an advantage of this system, as it could aid in learning more about specific particle subpopulations and to address the ability of phase state modulating proteins to act on these (e.g. ability to revert solid-like species back to liquid/gel-like phases).

XEE appear to be a valuable new addition to the current set of systems for studying biomolecular condensates. A main limitation of the XEE is that the extracts have a limited ‘lifespan’ and their composition changes over time. Moreover, the composition and quality of XEE can vary from batch to batch, which can introduce variability in experimental results (88, 89). However, XEE do offer a simplified system that allows to focus on specific cellular processes on a biochemical level, without the complexity of a whole living organism. The composition of XEE is well-characterized and plenty of techniques for manipulating and modifying these extracts have been developed (90, 91). This allows for exploring the effects of specific proteins, RNAs and/or other cellular components, as well physiologically-relevant environmental changes, on the formation and properties of biomolecular condensates. Also, cellular studies require the use of fluorescent tags, which might alter the properties of the target protein, including the tendency to form biomolecular condensates, while the XEE system allows the use of dye chemistry instead.

### PhaseMetrics is equally applicable for the assessment of biomolecular condensates in cells

Our analysis of particle properties in live yeast cells included a comparison between the plugin-based and manual assessment of particle distributions. This revealed a remarkable overlap between the findings of both approaches. However, the manual-based method requires much more time, is more subjective to human error, and did not include an accompanying detailed assessment of particle properties.

Furthermore, the plugin-based approach allows for further stratification of specific particle subpopulations, and can be easily further tailored in case classification into different types of particle categories would be desired. Our assessment of Nup100FG particle features in the absence and presence of DNAJB6b revealed that the plugin is sensitive enough to detect changes in a minor particle subpopulation. This further supports the potential of PhaseMetrics to assist in the identification of novel phase state modulators in an unbiased, high-throughput manner.

### Strengths and weaknesses of PhaseMetrics for studying biomolecular condensates

Overall, the findings from the three systems that we tested here – *in vitro* with chemically defined buffers, XEE, and yeast– are consistent, yet certain systems appear to be particularly favourable for monitoring specific particle subpopulations. Importantly, the pipeline is equally capable to analyse particle properties both *in vitro* and *in vivo*. Moreover, we compared the results from the imaging-based read-out to conventional biochemical assays used to study phase separation, which revealed the results from each of the assays are in good agreement. The main benefit of the imaging-based analysis is that it allows for high-throughput, quantitative assessment at single particle level, and is more sensitive in picking up differences in specific, minor particle subpopulations, whereas most biochemical methods report on population averages. Moreover, large-scale quantitative assessments allow for a more honest representation of variability within datasets. In many instances, only a single snapshot is provided that may or may not accurately reflect the overall diversity in distribution within the sample (13). The variability in several parameters of the observed particles (e.g. large spread in size and intensity values), the existence of minor particle subpopulations that may be lost or hidden by population averages, as well as differences between different imaging setups, are illustrative of the importance of being able to report on sample distributions. The subcategorization of the particles into distinct subpopulations can not only be used to highlight the existence of minor subpopulations, but also proof useful in comparing and highlighting the overlap and differences amongst different setups and experimental conditions. Finally, due to the time-consuming nature of manual quantifications, scoring is typically limited in the number of features that can be assessed. Automated analyses allow for capturing many features, within for example *in vivo* systems allowing for single-cell resolution for hundreds of images at a much faster rate.

Whilst the pipeline shares overlapping features with several other publicly-available image analysis plugins (69–72), the main distinction is that PhaseMetrics was specifically developed with the aim in mind to report on features that are particularly relevant in the assessment of biomolecular condensate properties. The analysis parameters, including for example the measurements reporting on particle shape (circularity and perimeter) and spatial heterogeneity of the signal distribution within the particles (skewness), were specifically selected to aid in the detection of changing particle properties in response to environmental cues. Whilst some of the other available tools for image-based analysis offer great flexibility in parameter selection, and could also accommodate the inclusion of parameters specifically suited for detecting changing particle properties in response to the surrounding environment such as for example CellProfiler (70), this would first require familiarization with the software and/or require a certain degree of programming experience. Moreover, the building of a custom pipeline and/or subsequent tailoring of post-analysis processing steps (e.g. for computation of relevant derivative measurements) can be time-consuming. Due to the user-friendly ImageJ macro language-based design, and use of custom-designed excel macros and accompanying post-analysis measurement sheets, specifically tailored to the assessment of relevant properties for studying biomolecular condensates, the PhaseMetrics pipeline is easily accessible and requires minimal to no programming experience. While we here only showcase its application for Nup100FG particles, our systematic testing of PhaseMetrics’ performance on multiple experimental systems, ranging from *in vitro* systems to cellular applications, demonstrates that the pipeline is widely applicable and anticipated to work equally well for other condensate-forming proteins.

A limitation of the imaging-based characterization of particles is that the morphologies of liquid-like and gel-like condensates or more solid aggregated species may not always differ enough to allow categorisation in different groups. Moreover, the resolution limit of standard microscopy setups precludes the assessment of properties of smaller particles, and cannot detect the subtle heterogeneities in particle morphology, such as the fibre-like protrusions that can be observed with hsAFM (92). Hence, it should be noted that morphology alone cannot serve as a good proxy for the distinguishment between different phase states. Furthermore, the correlation between intensity and concentration within the dense phase will be impacted by quenching effects, so changes in intensity should be interpreted with caution.

In conclusion, the image analysis pipeline that we introduced here appears to be a valuable addition to the current repertoire of tools to study biomolecular condensates, in particular when used in combination with conventional biochemical assays. Together, they can proof useful in the quest of advancing our understanding of the morphological and physicochemical properties of different types of particles, and present a toolbox that can help to uncover new biology.

### Experimental procedures Internet Resources

https://forum.image.sc/ The Scientific Community Image Forum.

https://imagej.net/Welcome The official wiki homepage for the ImageJ Ecosystem, including ImageJ and Fiji.

https://imagej.net/Fiji/Downloads The wiki page for downloading Fiji.

### Protein purification and phase separation assays

The expression and purification of the yeast Nup100FG-domain and the molecular chaperone DNAJB6b were performed as described before (52). In short, the pSF350 expression plasmid, containing a His6-tag at the N-terminal end and a cysteine residue at the C-terminal end was used for expression of the Nup100FG fragment (aa1-580) and full-length DNAJB6b (**Table 2**). For cell lysis and protein purification, a 100 mM Tris-HCl, 2 M guanidine-HCl, pH 8.0 buffer was used. For labelling, the Nup100FG fragment was incubated with fluorescein-5-maleimide (5MF, Thermo-Scientific, 62245) and DNAJB6b with Alexa fluor^TM^ 594 C5-maleimide (AF594, Alexa fluor^TM^ 594 C5-maleimide, Thermo-Scientific, A10256). Proteins were concentrated using Vivaspin Protein Concentrator spin columns (Vivaspin 10/30 kDa MWCO Polyethersulfone, Cytiva, 28-9322-47, 28-9322-48) and stored at −80 °C at a final concentration of 100 μM in 100 mM Tris-HCl 2 M guanidine-HCl, pH 8.0 buffer, supplemented with 10% glycerol. To reduce the impact of the fluorescent labels on the protein behaviour, the fluorescein-5-maleimide-labeled Nup100FG (Nup100FG-5MF) and Alexa fluor^TM^ 594 C5-maleimide-labelled DNAJB6b (DNAJB6b-AF594) were mixed with unlabelled proteins at 1:5 and 1:20 ratios, respectively.

**Table 2.** Plasmids.

| Designation | Source or reference | Additional information |
| --- | --- | --- |
| pSF350-Nup100FG (plasmid) | PMID: 36302971 | His10-TEV-Nup100FG<sub>1-580</sub>-Cys, IPTG inducible expression in *E. coli*, Amp<sup>R</sup> |
| pSF350-DNAJB6b (plasmid) | PMID: 36302971 | His10-TEV-DNAJB6b-Cys, IPTG inducible expression in *E. coli*, Amp<sup>R</sup> |
| pGAL1-eGFP-Nup100FG (plasmid) | This study | GAL1 promoter, N-term eGFP-Nup100FG<sub>1-580</sub>, His3+ Adapted from pACM021(93). PMID: 21659568 |
| pGAL1-mCherry (plasmid) | This study | GAL1 promoter, mCherry, Ura3+ Adapted from pACM063-mCh-L(180)-TM(94). PMID: 23357007 |
| pGAL1-mcherry-DNAJB6b (plasmid) | This study | GAL1 promoter, N-term mCherry-tag-DNAJB6b, Ura3+ Adapted from pACM063-mCh-L(180)-TM(94). PMID: 23357007 |

To assay phase separation behaviour, the purified proteins were diluted to a protein concentration of 3 μM in assay buffer. The assay buffer comprises 50 mM Tris-HCl pH8.0 and, as indicated, varying NaCl concentrations (0, 150mM, 500mM, 1M, 2M and 3M), supplemented with either no crowding agent or 10% w/v of polyethylene glycol 3350 (PEG3350; Sigma-Aldrich, P4338-2KG) for one hour at room temperature.

To assess whether PEG would be excluded from the particles, as would be expected from an inert crowding agent, we made use of PEG-rhodamine (mPEG-Rhodamine, MW 2k or 5k; Creative PEGworks, PSB-2265/2264). A 50 wt% PEG-rhodamine stock solution was prepared by mixing 495 mg/mL PEG (average molecular weight of 3.35k) with 5 mg/mL mPEG-Rhodamine (average molecular weight of 2 or 5k); the stock solution was diluted out into assay buffer to a final concentration of 10% PEG. Unlabelled Nup100FG was diluted out to a protein concentration of 3 μM into the 10% mPEG-rhodamine containing assay buffer and left to phase separate for one hour at RT.

To assess hexanediol (HD)-sensitivity of the Nup100FG particles, Nup100FG particles were either formed in the presence of 5% 1,6-HD or exposed for 10 min to 5% 1,6-HD after one hour of particle formation. To assess the effect of DNAJB6b on Nup100FG particle properties, 3 μM of Nup100FG-5MF and DNAJB6b-AF594 were mixed (1:1 molar ratio), and left to phase separate for one hour at room temperature. For the Nup100FG:DNAJB6b time course experiments, Nup100FG particle formation was followed over a three and a half-hour timeframe, with 30 min time-intervals. For the first replicate experiment, particle formation was assessed only up to three hours.

### Sample preparation for imaging of particles

Two different setups were used for studying particle formation using microscopy (**Figure 8A,B**). In the first setup, the proteins are left to phase separate in a low-protein-binding tube containing assay buffer. Right before imaging a 2 μL sample is mounted on an untreated glass-slide with coverslip (glass-slide setup). In the second setup, the protein samples are directly pipetted onto the surface of a chambered coverslip (Ibidi, #80826) and resuspended in a droplet of 10 μL assay buffer, followed by microscopy analysis (droplet-in-chamber setup). The main benefits of the glass-slide setup are its ease of use, and that the results are typically more homogeneous within the sample and more consistent between experiments. Yet, measurement of the soluble fraction does not appear to be accurate in the glass-slide setup, as it does not align with results from sedimentation assays, possibly due to protein-glass surface interactions. The main benefits of the droplet-in-chamber setup are that it allows for assessment of both the soluble and dense phases, and allows for studying real-time particle dynamics. Careful attention should be given to properly mix the sample to minimize the inhomogeneous distribution of particles throughout the droplet as much as possible (see **Figure S5**). This is particularly important when multiple proteins are mixed at variable time intervals. The main limitation of the droplet-in-chamber setup is that due to larger variability in the particle population within the sample, the thresholding can be more complicated. Whenever the accurate thresholding of particles requires time-consuming manual interventions, one may decide to only use the droplet-in-chamber setup for the extraction of the soluble fraction measurement. In this scenario, a thresholding method can be selected (see below for further details) that makes an overestimation of the particle population, to exclude particles from being included in the soluble fraction measurement.

### Xenopus laevis egg extracts (XEE)

Cytostatic factor-arrested (CSF) extracts were prepared in the presence of CSF-XB buffer (100mM KCl, 0.1 mM CaCl_2_, 2mM MgCl_2_, 10mM K-HEPES, pH 7.7, 50mM sucrose, 5 mM EGTA, pH 7.7, 10 μg/mL leupeptin, chymostatin, pepstatin and cytochalasin) as described previously (95), except that no energy mix was added. CSF extracts were stored at −80 °C until use and were left to thaw for 5 min at 20°C with mild shaking (<300 rpm). Cycloheximide was added to a final concentration of 100 µg/mL, followed by the addition of CaCl_2_ to a final concentration of 0.8 mM. The XEE was carefully mixed using low-retention pipet tips and incubated for 50 min at 20°C with careful mixing by stirring with a pipet tip every 10 min. The addition of CaCl_2_ initiates the meiotically-arrested extract to progress into interphase. Cycloheximide then stably arrests the extract in interphase by blocking the synthesis of cell cycle regulators (e.g. cyclin B). Right before imaging, Nup100FG-5MF was added at a final concentration of 3 μM to the interphase-arrested extracts, and 2 μL samples were mounted on glass-slides with coverslip at the indicated time points. *Xenopus laevis* are housed in accordance with national animal welfare laws and reviewed by the Animal Ethics Committee of the Royal Dutch Academy of Arts and Sciences (KNAW) under a project license granted by the Central Committee Animal Experimentation (CCD) of the Dutch government and approved by the University of Groningen (IvD), with project license number AVD 10500202114408.

### Yeast cell culture

The yeast *Saccharomyces cerevisiae* strain BY4741 (haploid) was used in this study. Yeast cells were grown at 30 °C for one day in selective drop-out (SD) medium containing 2% glucose (w/v) and then cultured for one day in medium containing 2% raffinose (w/v) as carbon source. On the day of the experiment, eGFP-Nup100FG (pGAL1-eGFP-Nup100FG-His3), and mCherry (pGAL1-mCherry) or mCherry-DNAJB6b (pGAL1-mCherry-DNAJB6b) (**Table 2**) expression was induced in exponentially growing cells with 1% (w/v) D-galactose for 2, 4 or 6 hours as indicated. 1 mL of culture was concentrated (1200x*g* for 1-2 min) to 50 μL, of which 2 μL was mounted on a glass slide with coverslip.

### Fluorescence microscopy

Microscopy was performed in a temperature-controlled environment at either 20 °C (*in vitro* and XEE) or 30 °C (yeast cells) using a DeltaVision Deconvolution Microscope (Applied Precision (GE), Issaquah, WA, USA), using an Olympus UPLS Apo 100x oil-immersion objective (NA 1.4) and InsightSSITM Solid State Illumination using the FITC 525/48 and A594 625/45 filter sets. Detection was done with either a CoolSNAP HQ2 or EDGE sCMOS5.5 camera. Fluorescence images were acquired with 30 Z-slices of 0.2 μm, and reference images were acquired at the middle of the sample using polarized light. Images were deconvolved using softWoRx software (Cytiva) and processed using the open-source software FIJI/ImageJ, making use of a custom-designed plugin presented here, called PhaseMetrics, for semi-automated analysis of particle properties.

### Image analysis

#### Manual assessment

For manual assessment of eGFP-Nup100FG particles in yeast cells, 150-250 cells were analysed for each independent replicate using FIJI. To enhance visualization of all particles, a maximum intensity projection of the eGFP channel was created, after which the brightness of the eGFP channel was standardly set to a range of 0-15000. This projection was subsequently merged with the brightfield channel to quantify the total number of particles per cell. Cells that lie on the image border were excluded from analysis. The particles were categorized into different subpopulations based on morphological appearance, including condensates (0, 1, 2, ≥3 condensates/cell), amorphous aggregates (AA) and amyloid-like bodies (AB), and their occurrence was evaluated on a per cell basis. The AB category is characterized by a fibre-like shape, the AA category by an irregularly-shaped structure that does not qualify as condensate nor as AB, and the condensates are characterized by a circular appearance. In the manual analysis, whenever a cell contains an AA or AB, they are grouped into these categories, regardless of the additional presence of condensates within these cells. In the automated analysis these cells are further stratified. This difference in analysis explains the small differences between the results of the manual and automated analysis of the particle distributions.

#### Plugin design and implementation

The PhaseMetrics plugin (**Figure 9**) was developed for the FIJI distribution of ImageJ. The pipeline is written in ImageJ macro language, making it easily accessible to all users, regardless of their programming experience. It makes use of both custom-designed and built-in ImageJ functions and plugins, and does not require any additional downloads for functionality, besides the activation a few update sites for non-standard plugins within FIJI. The plugin can be run using either the simple graphical user interface, or as command-line interface. Upon initiation, a graphical interface will appear that will request the user to select the desired settings for a number of parameters as defined below (**Table 3**). Prior to running the plugin for the first time, the user needs to manually define the desired settings and filters for thresholding. For this, the accompanying step-by-step PhaseMetrics instruction manual can be consulted. Depending on whether a single- or multi-channel image is to be analysed, two different modules can be run. The multi-channel module will analyse particles in both channels and includes a custom-designed object-based colocalization feature. Two different variants of the script were created, one specifically targeted at the analysis of *in vitro* experiments (PhaseMetrics_invitro), and one specifically targeted at the analysis of cellular experiments (PhaseMetrics_cellular). The main difference between these two variants is the type of colocalization analyses to be performed. Moreover, PhaseMetrics_cellular has been adjusted to allow for the analysis of three channels, whereas PhaseMetrics_invitro can only analyse up to two channels. The status of the analysis can be monitored in the log file, specifying the analysis step and image being processed. A message will appear at the end to indicate the analysis has finished. Output folders and results files will be automatically generated in the process. It is important to make use of a file directory that is fully accessible for proper export of the results files. Below, a more thorough description of each of the listed elements, as well as general guidelines on how to use this pipeline are given.

**Fig. 8.**
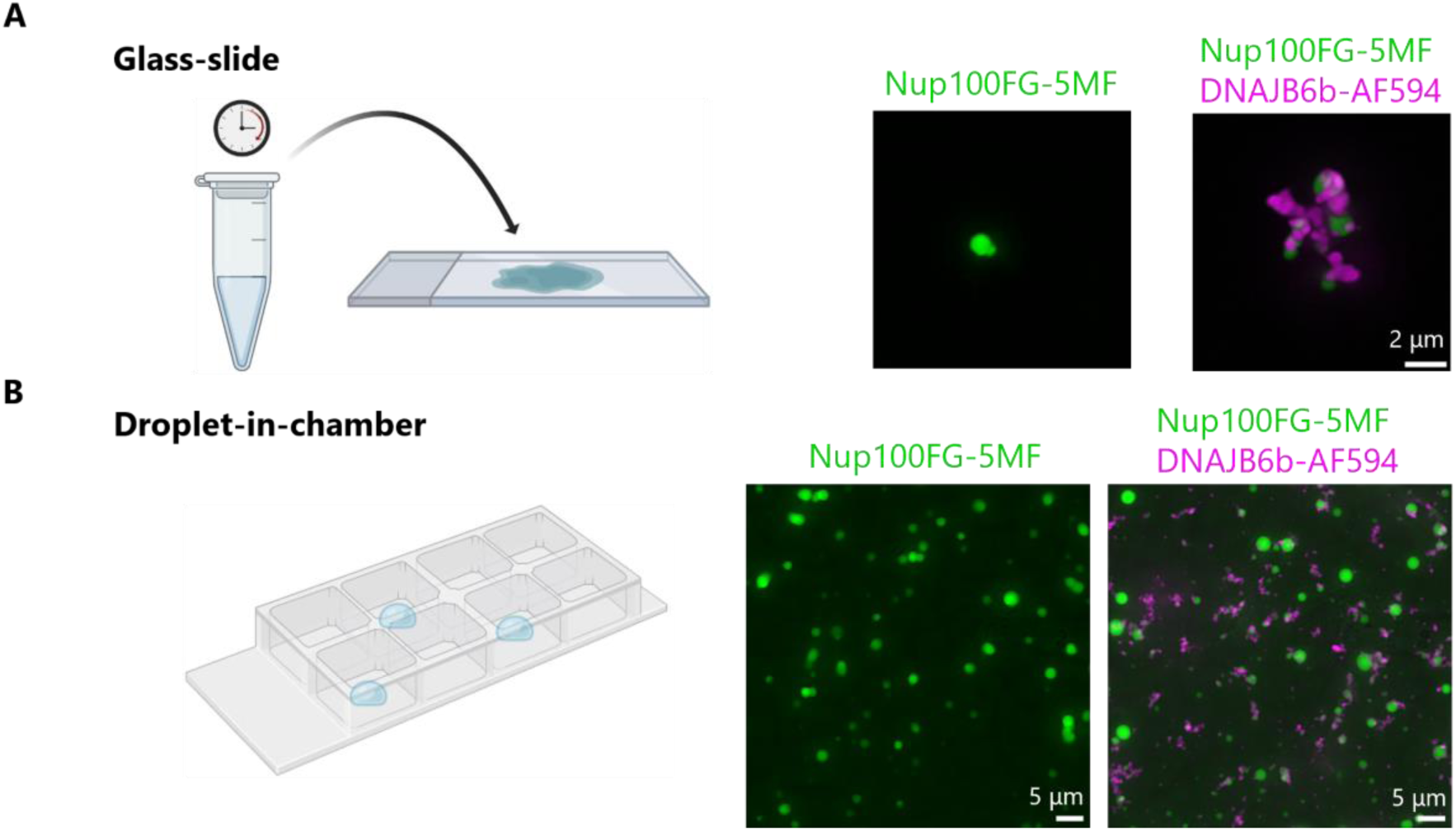
Overview of different types of particles and imaging setups to study particle formation. **(A,B)** Imaging setups to study particle formation. **(A)** In the glass-slide setup, proteins are left to phase separate for a pre-defined period of time in low-protein binding tubes, and mounted on an untreated glass-slide with coverslip right before imaging. Scale bar, 2 µm. **(B)** In the droplet-in-chamber setup, proteins are directly pipetted onto the surface of a chambered coverslip and resuspended in a droplet of assay buffer, followed by microscopy analysis. Scale bar, 5 µm.

**Fig. 9.**
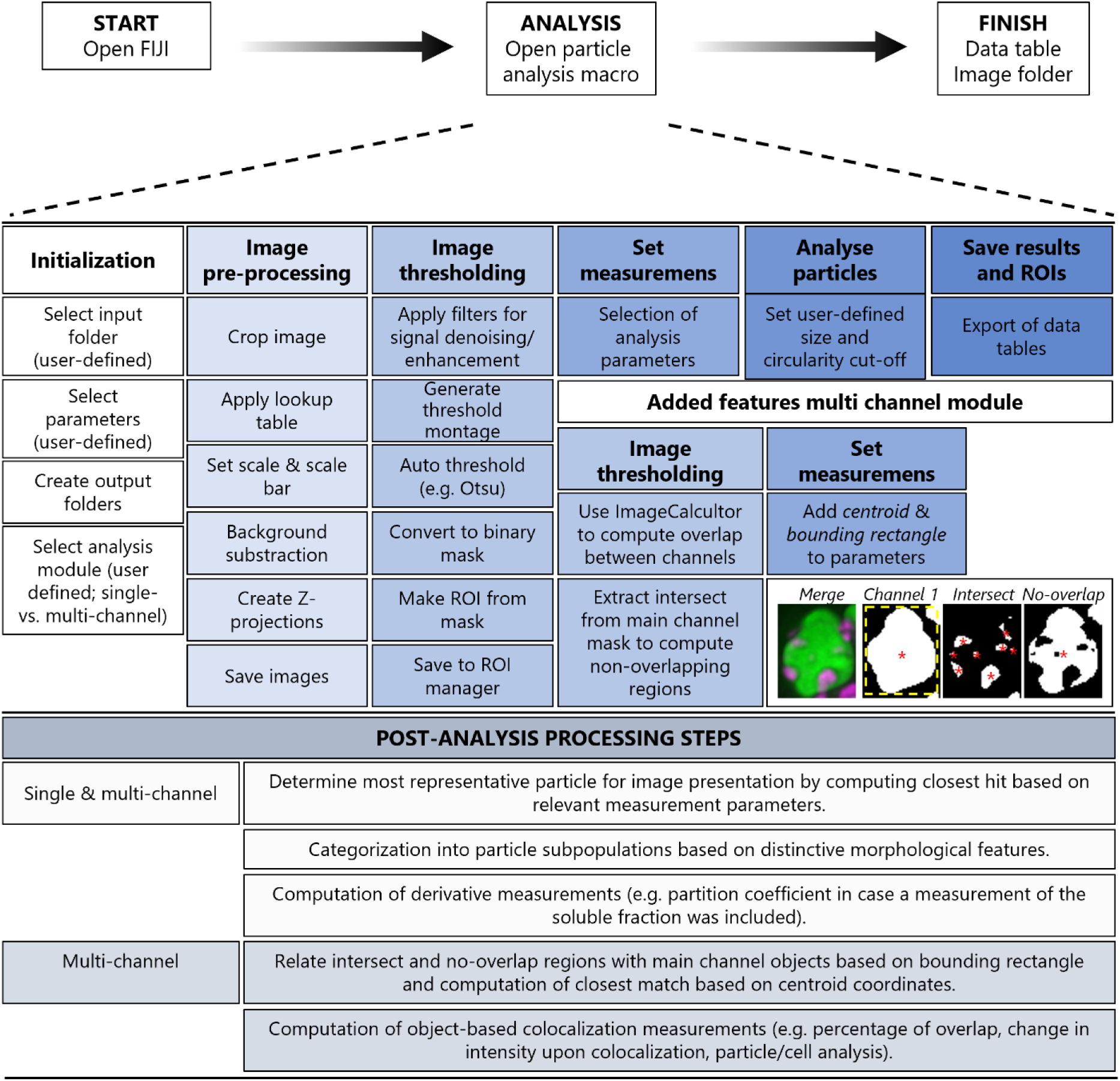
PhaseMetrics image analysis pipeline for quantitative assessment of biomolecular condensates. Plugin design with single channel module, and multi-channel module with object-based colocalization functionality. The flowchart describes the sequence of steps performed by the pipeline to extract multiple parameters from fluorescence microscopy images and allow for the analysis of biomolecular condensate properties. The yellow dashed box represents the bounding rectangle, red stars represent the centroids.

**Table 3.** List of parameters required for initiation, particle segmentation and analysis.

| Parameter | Description | Notes |
| --- | --- | --- |
| Input data | Directory with images to be analysed. | - |
| Imagetype | Requests for specification of file extension for images to be analysed. | Compatible with any file format that can be recognized by FIJI/ImageJ. Standardly set to R3D_D3D.dv file format. |
| Channels | Displays a dialog box to specify whether multi-channel module should be activated. | Will initiate the multi-channel module if user clicks "Yes" (1), the single-channel module will be initiated if user clicks "No" (0), and macro will exit if the user clicks "Cancel". |
| ImageCrop | Specify size dimensions (x,y, height and width) for setting image boundaries. | Leave blank in case no issues are encountered with particles locating at the edge of the imaging field, otherwise this setting can be used to crop off blurred edges to filter out particles that are partially out of frame. |
| ColorLUT_C# | Assigns desired colour to specified channel using ImageJ lookup tables. | - |
| Background subtraction_C# | Threshold value used to remove background signal in specified channel. | Uses built-in rolling-ball method to correct for uneven illumination. Areas of grey values large enough for a ball with the indicated radius to pass through are removed, while low intensity fine details are preserved. Radius should be set to at least the size of the largest object not being part of the background. |
| Filters_C# | Checkboxes for the selection of filters for noise reduction and/or signal enhancement. | *Median*: Reduces noise by replacing each pixel with median of itself and neighbouring pixel values within specified radius. Maintains borders while smoothening image.<br>*Mean*: Reduces noise by replacing each pixel with mean of itself and neighbouring pixel values within specified radius.<br>*Gaussian Blur*: Similar to smoothing filters, but instead replaces pixel value with a value proportional to a normal distribution of its neighbouring pixels (within specified radius).<br>*Unsharp mask*: Sharpens and enhances contrast by subtracting a blurred copy or unsharp mask from original image and rescales image to restore original contrast of large, low-frequency structures.<br>*Maximum*: Also known as dilation filter. Performs greyscale dilation by replacing each pixel in image with largest pixel value in that pixel's neighbourhood (within specified radius). Can be used to identify pixels with a greater value than all its neighbours, representing a local maximum.<br>*Minimum*: Also known as erosion filter. Performs greyscale erosion by replacing each pixel in the image with smallest pixel value in that pixel's neighbourhood (within specified radius). Can be used to identify pixels with a lower value than all its neighbours, representing a local minimum.<br>*Tophat*: Extracts small elements and details from original image, leading to homogenization of background. Can be used to enhance segmentation of bright structures. |
| Thresholding method_C# | Selection of thresholding algorithm for detection of ROIs in specified channel. | Offers full selection of built-in histogram-derived thresholding methods for image segmentation. Will result in creation of masked image for object identification. The threshold montage function of FIJI can be used to evaluate all available built-in thresholding methods. |
| Area cutoff_C# | The minimum and maximum allowed sizes in $\mu\text{m}$ to allow for careful detection of particles in specified channel. | Decrease to detect smaller species, increase to avoid noise detection. |

#### Requirements

Both single and multi-channel 3D Z-stacked images can be analysed using the analysis pipeline, for which ideally the Bio-Formats Windowless Importer should be used (96). The Windowless Importer can be selected for within ImageJ’s Plugin menu. When this option is selected, all subsequent image files will be imported using the last-used settings for a file of the same format. Additionally, the Excel Macro Extensions for ImageJ/Fiji are required, which can be installed by activating the update site *Excel Functions* in the *Help>Update* section of Fiji. A command is integrated into the plugin that upon initiation installs the extensions and makes them accessible via the ImageJ macro language script editor.

#### Plugin interfaces

As specified above, the plugin can be run using either a simple graphical user interface or as command-line interface. When opting for the simpler graphical user interface, the first step is to download the plugin into the FIJI plugins folder, and install it using the “Install…”option under the Plugins menu in the ImageJ toolbar. The plugin will subsequently appear in the Plugins menu. Alternatively, the script can simply be dropped onto the ImageJ toolbar and opened in the macro editor as a command-line interface, from which the “Run” button can be selected. This interface allows for more freedom in parameter selection and adaptation of folder and output file names.

#### Analysis initiation

**Table 3** lists the parameters that are required for the initiation of the analysis pipeline. Upon initiation, a dialog-box will appear requesting user input on the desired settings. First of all, an input folder has to be specified. This folder can contain different image file types, and hence one needs to specify the specific image type to be considered for analysis. All images within the same folder should be named systematically, and ideally different conditions should be split into separate folders to enhance overall processing speed, and allow for easy comparison between conditions. In case multi-channel images are being analysed, the user should assign each of the channels appropriately to correctly detect objects in each of the channels. Depending on whether the user is interested in analysing single or multi-channel images, one out of two different modules can be selected. The multi-channel module has the additive functionality to run an object-based colocalization test. The multi-channel module can be initiated by clicking “Yes” for “Multichannel?” in the dialog box during setup. In case the single-channel module is selected, all parameter requests for the other channel(s) can be ignored during setup. Additionally, the user is requested to indicate whether a measurement of the soluble fraction is desired. Depending on whether the PhaseMetrics_invitro or PhaseMetrics_cellular variant is used, this measurement is either 1) computed by extracting the particle mask from the original image, 2) or by extracting the particle mask from the cell mask, and measuring the background fluorescence in the remaining area, respectively. By subsequently computing the ratio of the dense (mean particle fluorescence) over light (mean background fluorescence) phase concentration, this measurement can be used to compute the extent of partitioning of biomolecules into particles, known as the particle partition coefficient (63, 97).

Once the appropriate settings have been defined, batch processing of all images in the specified folder will be initiated using the selected settings. The ImageCrop parameter requests for user input on what size dimensions should be used to set the image boundaries, which may be necessary when multiple objects are located at the edge of the imaging field to exclude detection of particles that are partially out of frame. The ColorLUT parameter can be used to assign the desired lookup tables to each of the channels to be analysed. The background subtraction parameter requests for user input to specify the “rolling ball” radius for background subtraction (see **Table 3** for further specification). This method allows for removal of large spatial variations in background intensities (98). The Filters parameter can be used to specify the desired filtering steps to be applied for noise reduction and/or signal enhancement prior to thresholding. These filters can both assist in reducing intensity variations and to amplify weak signals that would otherwise be missed during thresholding (see **Table 3** for further specification). **Figure S6** represents illustrative examples of how specific filters can be used for improved detection of objects displaying large intensity variations. Similar filters for noise reduction and signal amplification can be useful in the assessment of particle properties in cells, where the signal may be more noisy and more complex mixtures of object types may arise, resulting in great intensity variations that can complicate thresholding. The thresholding-method parameter requests for user input on what thresholding algorithm to use for object segmentation. In case the multi-channel module is used, thresholding will be applied to each of the individual channels to create object masks. For the *in vitro* experiments, these masks will subsequently be used for determining the intersected and no-overlap regions between the particles observed in both channels. For the cellular experiments, these masks will be used to determine which particles overlap with a cell mask, and when a third channel is included (e.g. for the detection of a phase state modulating protein) assess whether the particles observed in both channels colocalize. Finally, the area cut-off parameter can be used to specify the minimally and maximally allowed size for fateful detection of particles.

#### Measurements

The most important measurements that will be automatically extracted from all objects, along with a short description of their relevance towards evaluating the properties of particles, are specified in **Table 4**. They include a measurement of its size (area), and mean intensity, as well as parameters that are descriptive of its shape (circularity and perimeter) and a parameter that captures the spatial heterogeneity (skewness). For co-localization purposes, two additional measurements are included namely the “centroid” and “bounding rectangle” measurements. These represent the x and y coordinates for the centre point of the detected objects and for the smallest rectangle enclosing the objects respectively, which is used to relate objects between the different channels. For two objects to be considered co-localized, XY coordinates of the specified intersected and no-overlap regions should lie within their bounding rectangle and be near the coordinates of the “main” objects (explained in more detail below).

**Table 4.**
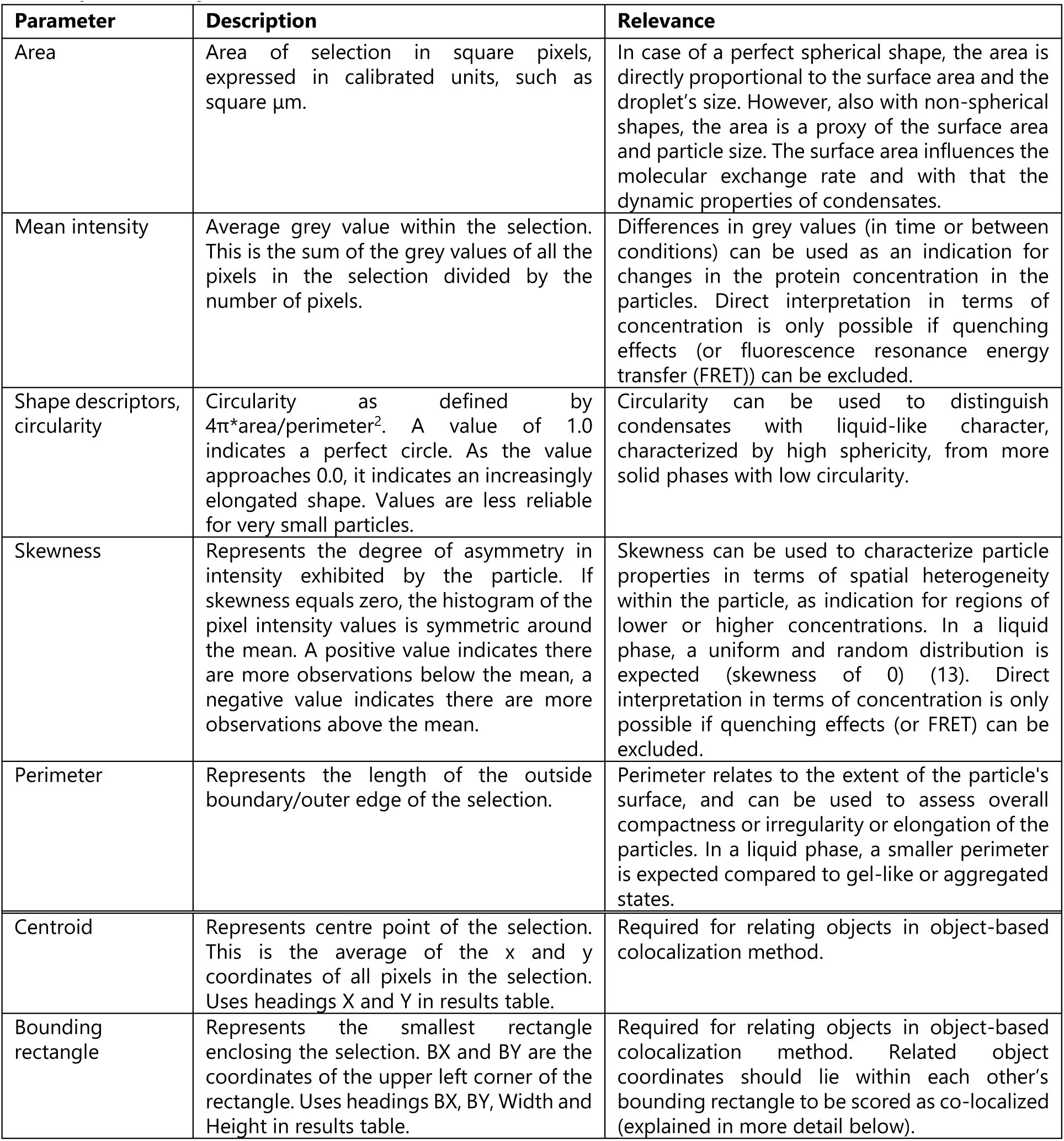
List of analysis measurements.

| Parameter | Description | Relevance |
| --- | --- | --- |
| Area | Area of selection in square pixels, expressed in calibrated units, such as square $\mu\text{m}$. | In case of a perfect spherical shape, the area is directly proportional to the surface area and the droplet's size. However, also with non-spherical shapes, the area is a proxy of the surface area and particle size. The surface area influences the molecular exchange rate and with that the dynamic properties of condensates. |
| Mean intensity | Average grey value within the selection. This is the sum of the grey values of all the pixels in the selection divided by the number of pixels. | Differences in grey values (in time or between conditions) can be used as an indication for changes in the protein concentration in the particles. Direct interpretation in terms of concentration is only possible if quenching effects (or fluorescence resonance energy transfer (FRET)) can be excluded. |
| Shape descriptors, circularity | Circularity as defined by $4\pi \cdot \text{area} / \text{perimeter}^2$. A value of 1.0 indicates a perfect circle. As the value approaches 0.0, it indicates an increasingly elongated shape. Values are less reliable for very small particles. | Circularity can be used to distinguish condensates with liquid-like character, characterized by high sphericity, from more solid phases with low circularity. |
| Skewness | Represents the degree of asymmetry in intensity exhibited by the particle. If skewness equals zero, the histogram of the pixel intensity values is symmetric around the mean. A positive value indicates there are more observations below the mean, a negative value indicates there are more observations above the mean. | Skewness can be used to characterize particle properties in terms of spatial heterogeneity within the particle, as indication for regions of lower or higher concentrations. In a liquid phase, a uniform and random distribution is expected (skewness of 0) (13). Direct interpretation in terms of concentration is only possible if quenching effects (or FRET) can be excluded. |
| Perimeter | Represents the length of the outside boundary/outer edge of the selection. | Perimeter relates to the extent of the particle's surface, and can be used to assess overall compactness or irregularity or elongation of the particles. In a liquid phase, a smaller perimeter is expected compared to gel-like or aggregated states. |
| Centroid | Represents centre point of the selection. This is the average of the x and y coordinates of all pixels in the selection. Uses headings X and Y in results table. | Required for relating objects in object-based colocalization method. |
| Bounding rectangle | Represents the smallest rectangle enclosing the selection. BX and BY are the coordinates of the upper left corner of the rectangle. Uses headings BX, BY, Width and Height in results table. | Required for relating objects in object-based colocalization method. Related object coordinates should lie within each other's bounding rectangle to be scored as co-localized (explained in more detail below). |

#### Output folders

The plugin will automatically create new folders within the user-specified directory to save the analysis output, including a folder containing the max intensity (Max) and sum of slices (Sum) Z-projection images as .tiff files. The maximum intensity projection is used for creating the segmented image for object detection after which the object masks are re-directed to the sum of slices projection to extract the desired measurements. Additionally, a Montage folder is created for saving the threshold montage images, which can be used for reference to verify threshold settings. The results folder will contain a single excel file with the quantifications of the specified measurements for each of the detected objects in all channels analysed. In case the object-based colocalization modules were selected, the results sheet will also contain tabs with all extracted measurements from 1) the intersected and no-overlap regions for the *in vitro* datasets, and 2) particles colocalizing with a cell mask, soluble measurements, and the intersected regions between the particles and a colocalizing protein (e.g. phase state modulating protein). Finally, a ROIs folder will be created for the .zip files comprising the ROIs for all detected objects for each image.

#### Post-analysis processing steps

Once the plugin-based analysis has been completed, the accompanying macro-containing excel sheet (ExcelMacrosWorkbook.xlsm) can be used for post-analysis processing of the exported results table. This sheet contains four different modules, including a: 1) ‘Particle selection & Particle distribution’ macro, 2) ‘Computation partition coefficient’ macro, 3) Two subvariants of a ‘Colocalization analysis’ macro tailored for *in vitro* datasets, and 4) Three subvariants of a ‘Colocalization analysis’ macro tailored for cellular datasets. Details on processing steps and resulting output from these different macros are specified in the tables below the different modules, and instructions on how to use these macros are further detailed in the PhaseMetrics instruction manual. In short, the ‘Particle selection & Particle distribution’ macro will copy the required information from the results table into a pre-designed excel sheet (ReprParticleSelectionSheet_invitro.xlsx or ReprParticleSelectionSheet_cellular.xlsx) for the selection of the particle that best represents the complete dataset, incorporating the area, mean intensity, perimeter, circularity, and skewness measurements. Additionally, it will categorize the particle population into multiple subpopulations (condensates, amorphous aggregates (AA) and amyloid-like bodies (AB)), making use of pre-defined search criteria, and return the most representative particles for each of these subpopulations. The currently used search criteria were determined by manual scoring of ≥60 particles (∼20 particles for 3 replicates) for each of the specified subpopulations, based on their morphological properties, using the corresponding mean, minimum and maximum circularity, size, and perimeter values. These subpopulations and search criteria can be manually adjusted if desired. The worksheets used for the categorization currently permit for a maximum of three different subpopulations, but this can be further expanded if desired, in which the applied formulas for the current subcategorization can be used as reference. The ‘Computation partition coefficient’ macro will link the mean background fluorescence values to the particles detected in the corresponding images, and calculate the partition coefficient by dividing the dense by light phase concentration. This computation is also incorporated in the colocalization analyses when a measurement of the soluble fraction was included. The ‘Colocalization analysis’ macros make use of two pre-designed excel sheets, one to determine which objects are colocalized (RelatingObjectsSheet_invitro.xlsx or RelatingObjectsSheet_cellular.xlsx) and one for the subsequent colocalization measurements (ColocalizationSheet_invitro.xlsx or ColocalizationSheet_cellular.xlsx). Two objects are considered colocalized when the intersected region lies within the bounding rectangle of the main object. If this criterion is not met (e.g. because the centre point of the intersected region lies exactly at the edge of the colocalized object), the closest object will be computed based on the XY coordinates of the centre points (centroids) of both objects. This distance between the centroids of both objects may not exceed a pre-determined maximum distance cut-off, which has been manually determined for the analysed datasets and is standardly set to 1.99 μm. This cut-off can be easily adjusted in the accompanying RelatingObjectsSheets if desired. The main objects that belong to each of the intersected and/or no-overlap regions will subsequently be displayed in the ‘Relate’ column. This column will be pasted into the corresponding colocalization measurements sheet (ColocalizationSheet_invitro.xlsx or ColocalizationSheet_cellular.xlsx) to link corresponding objects to one another and separate non-colocalized objects from colocalized objects. Moreover, this sheet will automatically compute several colocalization-based measurements, which vary depending on the selected subvariants and whether an *in vitro* or cellular dataset is being analysed. The most-important read-outs from each of the different analyses will be summarized in the first ‘Summary’ tab. Upon initiation of the excel macros, a dialog box will appear requesting the desired excel sheets. Upon completion, each of the excel sheets used for the colocalization test will be automatically saved for later reference.

#### Cell lysis, sedimentation assay and immunoblotting

Budding yeast cells overexpressing eGFP-Nup100FG (and mCherry /mCherry-DNAJB6b) were collected by centrifugation, washed with PBS, and frozen in liquid nitrogen. Cell pellets were resuspended in HEPES buffer (50 mM HEPES pH 7.5, 100 mM NaCl, 2.5 mM MgCl_2_, 10 mM DTT and 10% Glycerol) supplemented with protease inhibitors (10 mM PMSF and cOmplete-EDTA protease inhibitor cocktail), and broken with glass bead using the Fast-Prep homogenizer (MP Biomedicals). The lysate was clarified by centrifugation at 3,000 x g for 3 min. Total protein concentration of whole cell extracts was quantified using Pierce^TM^ BCA Protein Assay Kit. Cell lysates were adjusted to the same concentration, preparing enough sample to perform the sedimentation assay, the immunoblot and the filter trap assay from the same sample.

To assess eGFP-Nup100FG protein levels by immunoblotting, equal amounts of cell lysates were diluted in 2X protein loading dye buffer and boiled for 30 min. For the sedimentation assays, 30 μL of each sample was centrifuged at 13,000 rpm for 10 min at room temperature, after which the soluble fraction (supernatant) was transferred to a new tube, while the insoluble fraction (pellet) was left in the same tube and resuspended in 30 μL HEPES buffer. Both fractions were mixed with 2X protein loading dye buffer and boiled for 30 min. Subsequently, samples were separated via SDS-PAGE (Stain free gels 10%, BioRad), and transferred to PVDF membranes. Membranes were blocked with 2.5% BSA in PBS-T (0.1%), incubated overnight with primary antibody monoclonal mouse anti-GFP (Santa Cruz, sc-9996) (1:2500), and 2h with secondary anti-mouse m-IgGk BP-HRP protein (Santa Cruz, sc-516102) (1:2500), and revealed with enhanced chemical luminescence (ECL) using the Chemidoc imaging system (BioRad). For quantification of the insoluble and soluble fractions, the corresponding band intensities were measured using FIJI, corrected by total protein loading (using the Stain Free signal), and then made relative to the average of the control.

#### Filter trap assay

For the filter trap assay (FTA), cells and protein samples were processed as explained above and, after equalizing the concentrations, samples were incubated with SDS (100 μL of cell extract with 1% SDS) for 10 min in shaking. FTAs were performed as described before (52). After this, membranes were processed and treated as explained for immunoblot experiments. For the quantification of the FTA, band intensities were measured using FIJI and expressed relative to the average intensity of the control.

#### Statistical analysis

Graph preparation and statistical tests were performed using Prism10 software (GraphPad). The number of biological and technical replicates are indicated for all experiments in the figure legends. A colour gradient was applied to highlight the individual datapoints belonging to each of the independent replicates, unless mentioned otherwise. Data was tested for normality using the Shapiro-Wilk and D’Agostino-Pearson tests. For normal distributed data, a parametric unpaired t-test (two groups) (Fig. 3G) or one-way ANOVA (three or more groups) (Fig. 3M) was performed. For non-normal distributed data, a non-parametric Mann Whitney test (two groups) (Fig. 3B-F,H; Fig. S3B; Fig. 6G; Fig. S4E, Fig. 9D-H) or Kruskall-Wallis test (three or more groups) (Fig. 3J-L,N; Fig. 4B-F, Fig. 5B-F, Fig. 6B-D, H-J; Fig. S4G; Fig. 7B-F, Fig. 8D-G) with Dunn’s multiple comparisons test was performed. For analysis of particle distributions, analysis of contingency by Chi-square was performed to identify differences between replicates. To test for statistical differences in the overall distribution of the particle subcategories between the different conditions, a Chi-square test (based on the raw numbers) was used. To test for differences between each of the specified particle subcategories, multiple parametric unpaired t-tests were used (based on the three replicates in percentages). To test for differences between band intensities following immunoblotting and detection using chemiluminescence reagents (ECL), parametric unpaired t-tests were used. Two-tailed P values; ∗ p=<0.05, ∗∗ p=<0.01, ∗∗∗ p=<0.001, **** p=<0.0001.

## Supporting information

This article contains supporting information (Supplementary files 1-3)

- Supporting Data 1: Supporting figures S1-S6.
- Supporting Data 2. Folder containing PhaseMetrics scripts:

`o` PhaseMetrics_cellular (FIJI script for cellular datasets)
`o` PhaseMetrics_invitro (FIJI script for *in vitro* datasets)
- Supporting Data 3. Folder containing PostAnalysisProcessingFiles:

`o` ReadMe (1aReadme.txt)
`o` Macro-containing excel sheet for post-analysis processing (1ExcelMacrosWorkbook.xlsm)
`o` Excel worksheets for identifying the most representative particle (2ReprParticleSelectionSheet_invitro.xls and 3ReprParticleSelectionSheet_cellular.xls)
`o` Excel worksheets for relating objects for object-based colocalization (4RelatingObjectsSheet_invitro.xls and 5RelatingObjectsSheet_cellular.xls)
`o` Excel worksheets for object-based colocalization measurements (6ColocalizationSheet_invitro.xls and 7ColocalizationSheet_cellular.xls)

## Data availability

All numerical data are contained within the manuscript. Raw imaging data and source code will be deposited into a publicly accessible repository, the location and identifier will be made available after acceptance. Example datasets and the accompanying instruction manual are available upon request. There are no exceptions or limitations to the sharing of data, materials, and software.

## Acknowledgement

We thank Amarins Blaauwbroek and Johan Zijlstra for practical assistance and Michael Chang and all members of the Veenhoff and Chang laboratories for their valuable suggestions. This work was financially supported by the Netherlands Organization of Scientific Research grant no. VI.C.192.031 and OCENW.GROOT.2019.068 to L.M.V.

## Author contributions

Tessa Bergsma: Conceptualization, Methodology, Software, Validation, Formal analysis, Investigation, Visualization, Writing. Anton Steen: Methodology, Writing-review, Investigation. Julia Kamenz: Methodology, Resources, Writing-review. Paola Gallardo: Methodology, Formal analysis, Investigation, Visualization, Writing-review and editing. Liesbeth Veenhoff: Conceptualization, Writing-review and editing, funding acquisition, supervision.

---

**Fig. S1.**
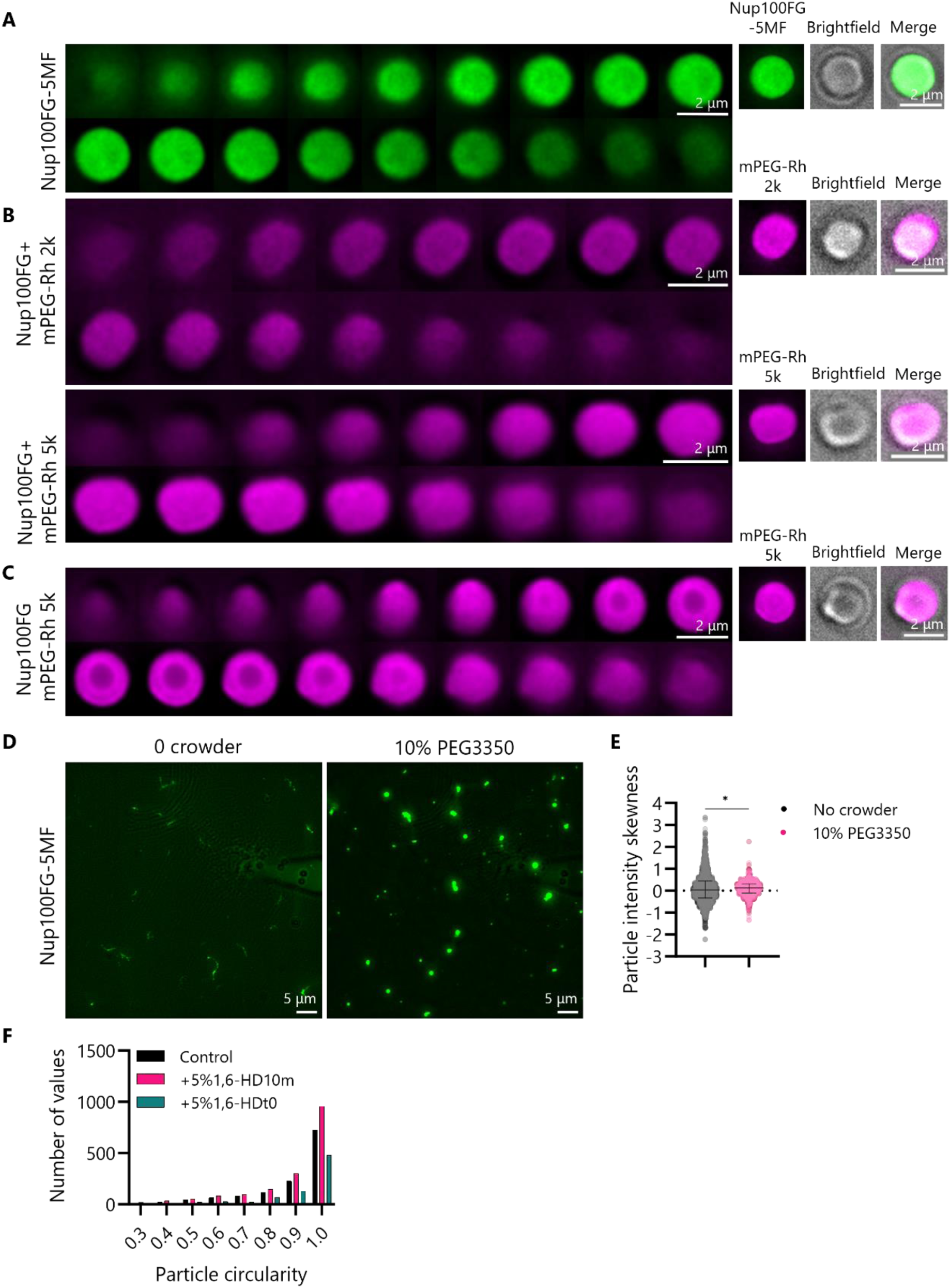
Characterization of the properties of Nup100FG particles formed in vitro. **(A,B)** Representative montage of 2 µm slices from Z-stack images of Nup100FG-5MF particles **(A)** and Nup100FG particles formed in the presence of mPEG-rhodamine (2k or 5k) **(B)**. Scale bar, 2 µm. **(C)** As in **(B)**, yet illustrating how mPEG-rhodamine 5k occasionally gets incorporated to a variable extent into the Nup100FG condensates. Scale bar, 2 µm. **(D)** Representative images of Nup100FG-5MF particles, formed in the absence or presence of 10% PEG3350 (1h) (droplet-in-chamber setup). Scale bar, 5 µm. **(E)** Intensity skewness of Nup100FG-5MF particles exemplified in **(D)**. Graphs show median ± interquartile range of ≥1000 particles per condition (n=3). *P<0.05. **(F)** Distribution plot showing the number of values for Nup100FG particle circularity, formed in the presence of 10% PEG3350 (1h), in the absence or upon exposure to 5% 1,6-hexanediol (n=2) (particles exemplified in **Figure 1I**).

**Fig. S2.**
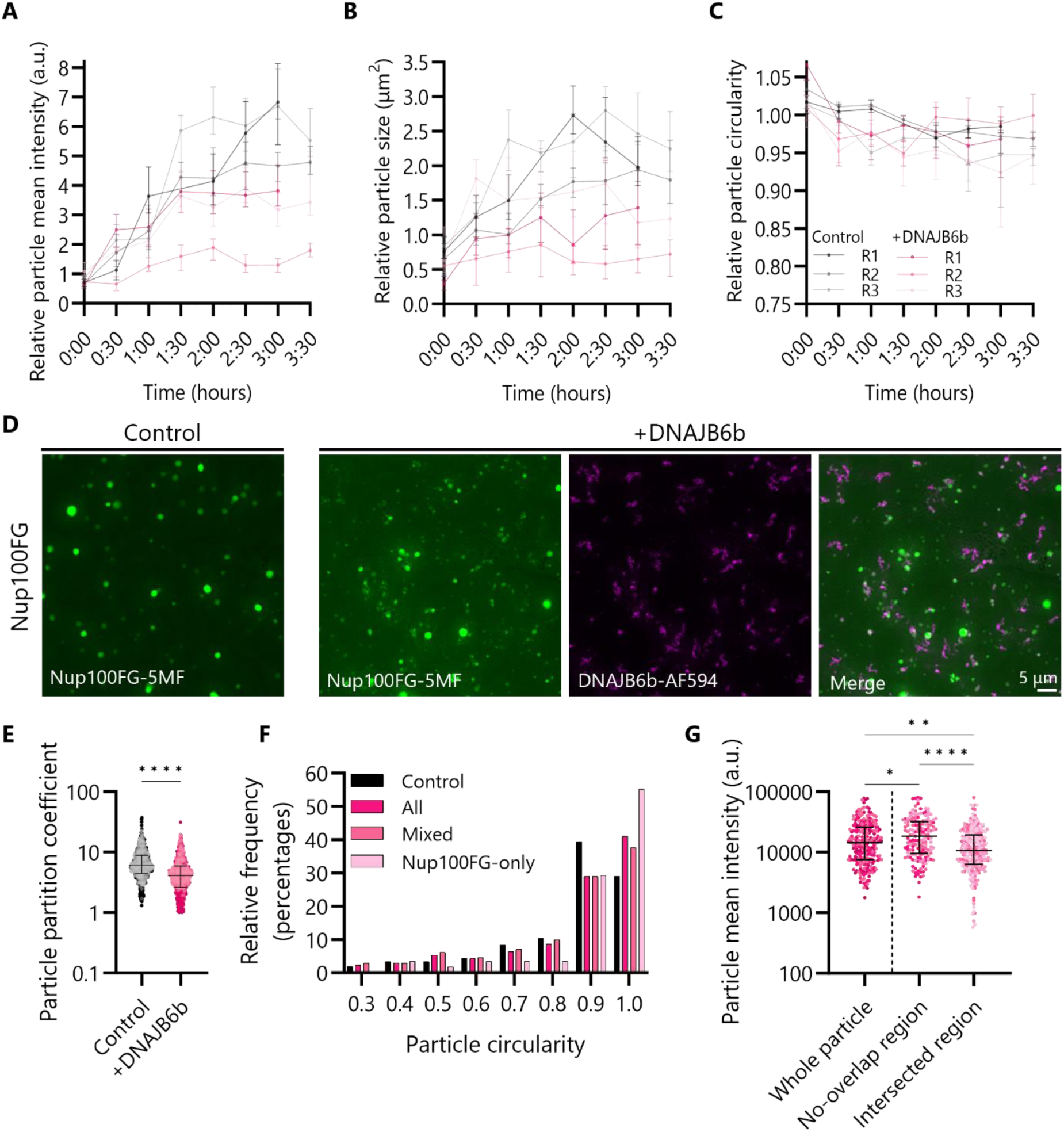
Quantitative assessment of changing Nup100FG particle properties in vitro in the presence of a phase state modulator. **(A-C)** Individual repetitions of the data in Figure 4A-D. Line graphs show the median ± 95% CI of 100 particles for each time point for each of the independent replicates. A colour gradient was used to highlight the data belonging to each of the independent replicates. **(D)** Representative images showing Nup100FG-5MF particles in the absence or presence of DNAJB6b-AF594 (1h) (molar ratio 1:1) (droplet-in-chamber setup). Scale bar, 5 µm. **(E)** Partition coefficient of Nup100FG-5MF±DNAJB6b protein mixtures exemplified in **(D)**. **(F)** Frequency distribution of circularity of Nup100FG-5MF particles formed in the absence or presence of DNAJB6b-AF594 (1h) (molar ratio 1:1), exemplified in (Figure 4F). Mixed: Nup100FG particles that are colocalized with a DNAJB6b particle, Nup100FG-only: Nup100FG particles that are not co-colocalized with a DNAJB6b particle. **(G)** Change in fluorescence intensity in different regions of Nup100FG-5MF particles colocalized with a DNAJB6-A594 particle, exemplified in (Figure 4F). Median ± interquartile range. *P<0.05, **P<0.01, ****P<0.0001.

**Fig. S3.**
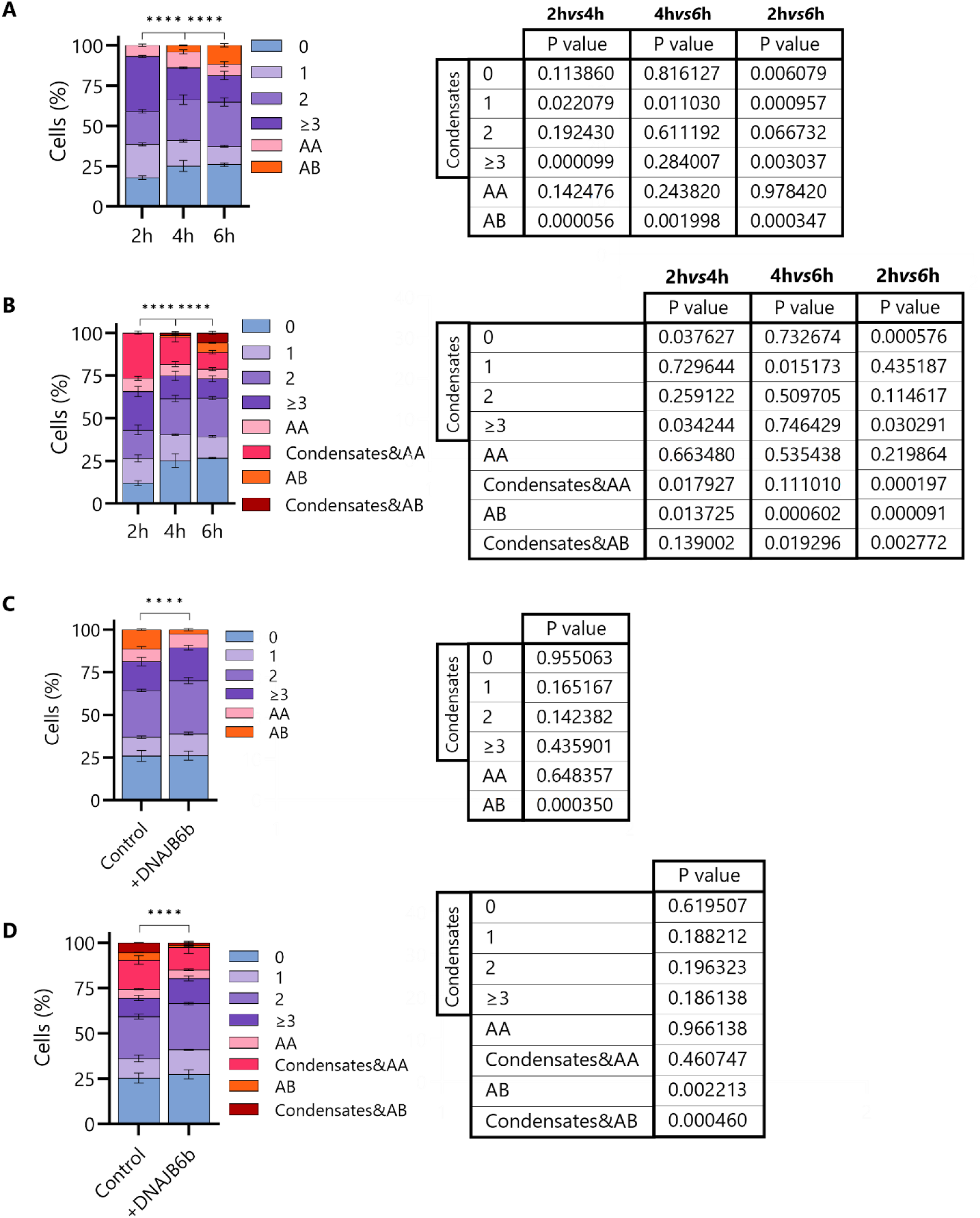
Comparison of manual and plugin-based quantification of particle distributions in yeast cells. **(A,B)** Distributions of eGFP-Nup100FG particles per cell, followed over time, using **(A)** manual and **(B)** plugin-based quantification. **(C,D)** Distributions of eGFP-Nup100FG particles per cell, in absence (mCherry as negative control) or presence of mCherry-DNAJB6b, using **(C)** manual and **(D)** plugin-based quantification. Graphs show mean ± SEM of ≥450 cells per condition (n=3). Grouped Chi-square < 0.0001. Tables represent multiple unpaired parametric t-tests of condensate distributions.

**Fig. S4.**
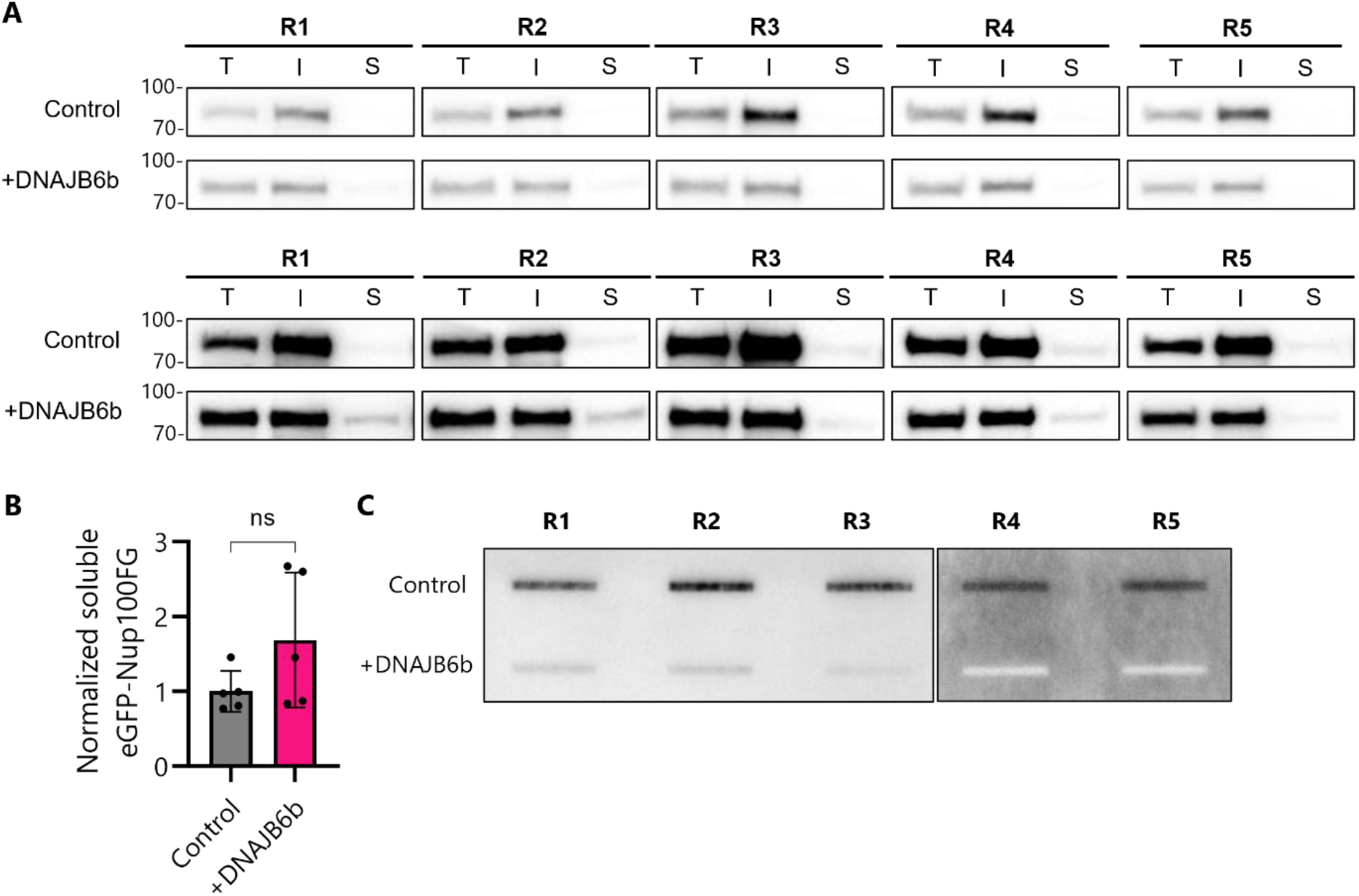
Individual repetitions of data in Fig. 9I-L. **(A)** Sedimentation assays to assess the soluble and insoluble fraction of eGFP-Nup100FG in the absence (mCherry as negative control) or presence of DNAJB6b. Bottom panel corresponds to the overexposed sedimentation assays, to highlight the soluble fraction of eGFP-Nup100FG. T: Total protein, I: Insoluble fraction, S: Soluble fraction. **(B)** Quantification of the soluble fraction of eGFP-Nup100FG. Represented band intensities are relative to the average intensity of the control. Mean ± SEM (n=5). **(C)** Filter trap assays to assess the SDS-insoluble fraction of eGFP-Nup100FG in the absence or presence of DNAJB6b.

**Fig. S5.**
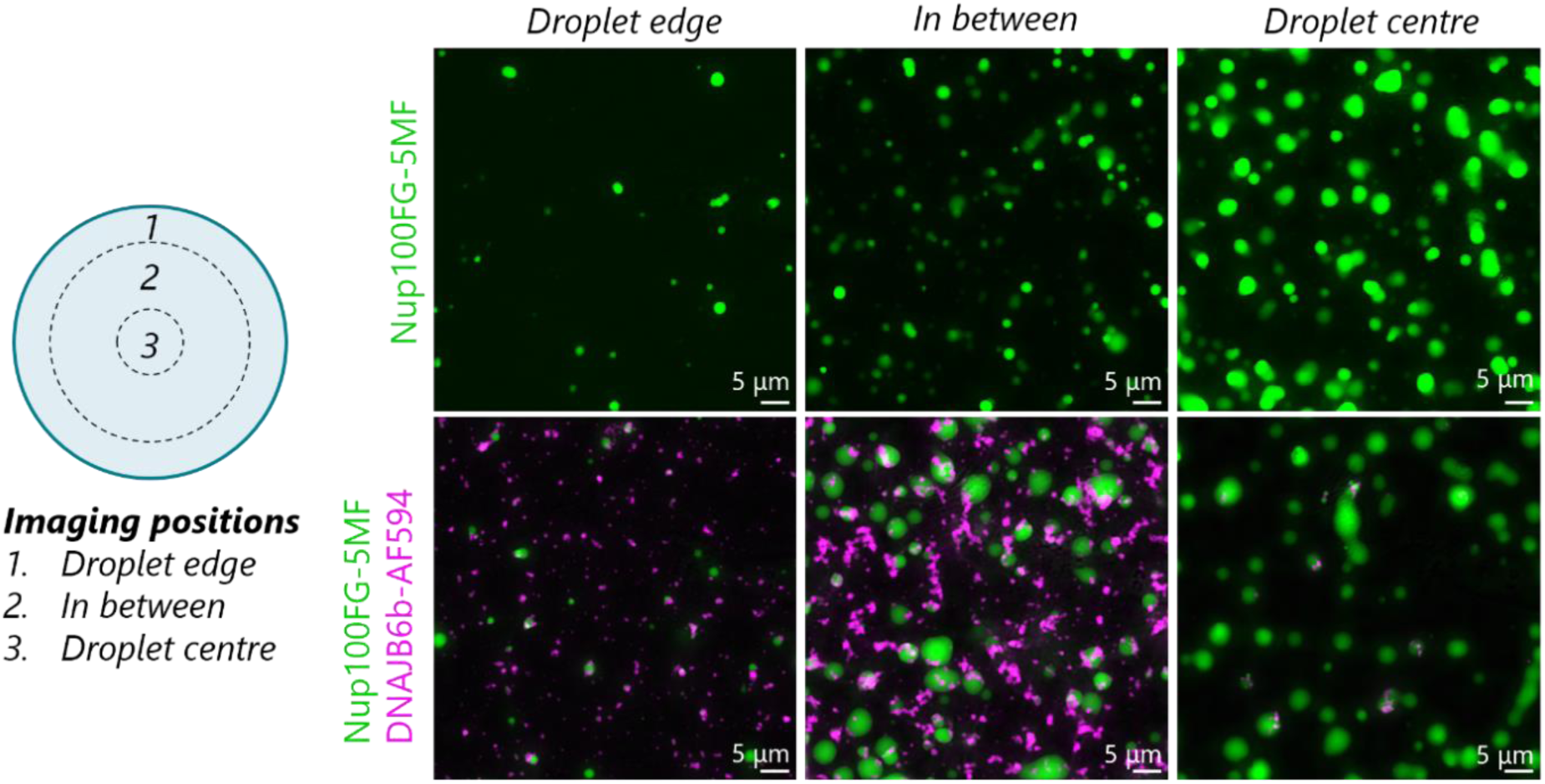
Inhomogeneous sample distribution droplet-in-chamber setup. The selected image frames illustrate how, if not properly mixed, this setup displays an inhomogeneous distribution throughout sample that complicates analysis of particle properties. Scale bar, 5 µm.

**Fig. S6.**
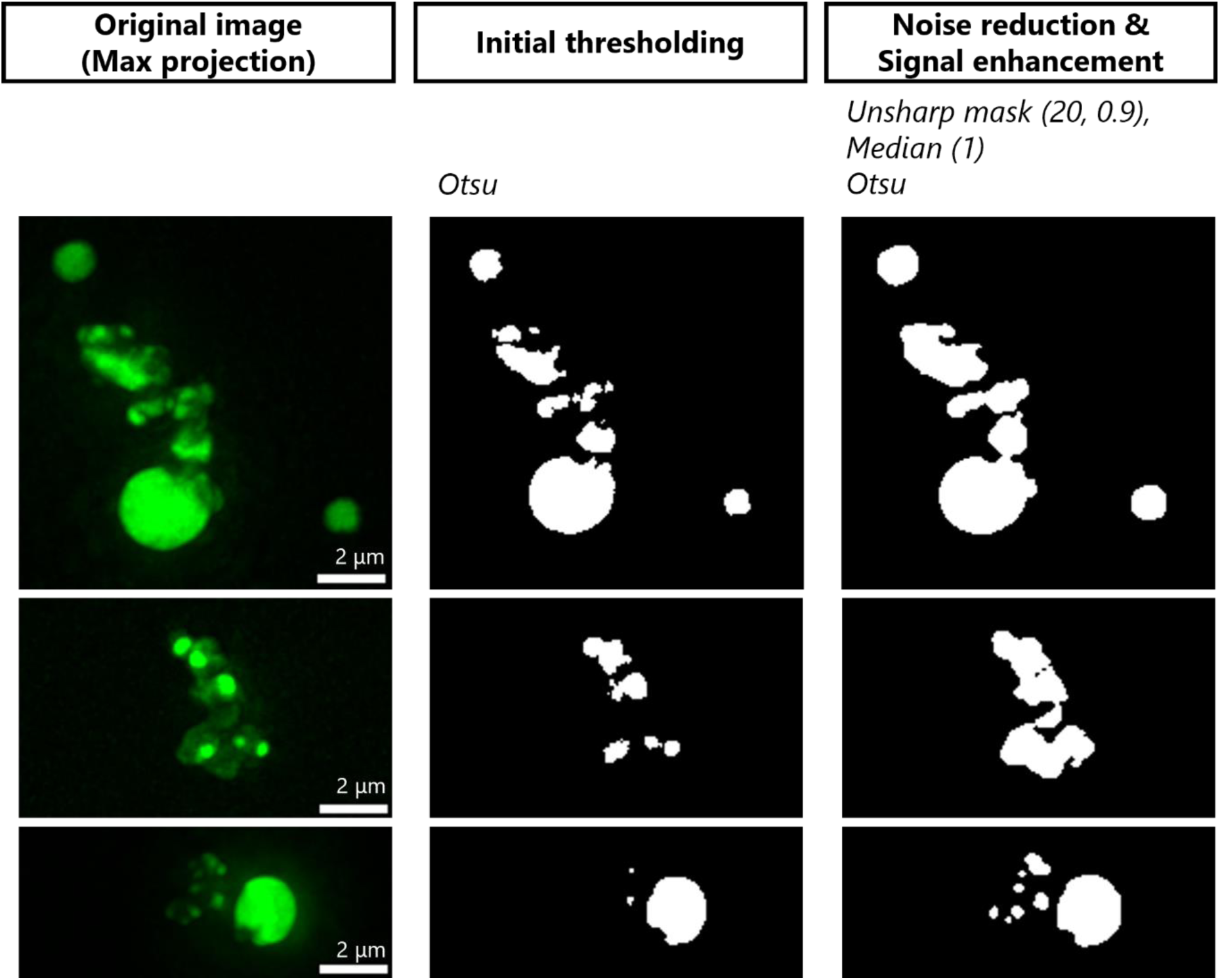
Examples of filtering options for improved object segmentation. Visual representation of the use of unsharp mask and median filters for noise reduction, sharpening and enhancing contrast for improved detection of objects displaying large intensity variations. Scale bar, 2 µm.

